# Genomic analyses of wild and cultivated bacanora agave (*Agave angustifolia* var. *pacifica*) reveal inbreeding, few signs of cultivation history and shallow population structure

**DOI:** 10.1101/2022.04.13.488215

**Authors:** Anastasia Klimova, Karen Y. Ruiz Mondragón, Francisco Molina Freaner, Erika Aguirre-Planter, Luis E. Eguiarte

**Affiliations:** Department of Evolutionary Ecology, Institute of Ecology, National Autonomous University of Mexico, Circuito Exterior s/n Annex to the Botanical Garden, 04510, Mexico City, Mexico; Department of Ecology of Biodiversity, Institute of Ecology, National Autonomous University of Mexico, Hermosillo, Sonora, 83250, Mexico

**Keywords:** gene flow, genomic resources, genomic variation, GBS, single nucleotide polymorphisms, spatial structure

## Abstract

Due to the recent increase in demand for agave-based beverages, many wild agave populations have experienced rapid decline and fragmentation; whereas cultivated plants are now managed at monocultural plantations, in some cases involving clonal propagation. We examined the relative effect of migration, genetic drift, natural selection and human activities on the genetic repertoire of *Agave angustifolia var. pacifica*, an agave used for bacanora (an alcoholic spirit similar to tequila) production *in northwestern* Mexico. We sampled 34 wild and cultivated sites and used over eleven thousand genome-wide SNPs. We found shallow genetic structure among wild samples, although, detected differentiation between coastal and inland sites. Surprisingly, no differentiation was found between cultivated and wild populations. Moreover, we detected moderate inbreeding (*F*_IS_ ∼ 0.13) and similar levels of genomic diversity in wild and cultivated agaves. Nevertheless, the cultivated plants had almost no private alleles and presented evidence of clonality. The overall low genetic structure in *A. angustifolia* var. *pacifica* is apparently the result of high dispersibility promoted by pollinators and possibility of clonal reproduction. Incipient cultivation history and reliance on wild seeds and plants are probably responsible for the observed patterns of high genetic connectivity and considerable diversity in cultivated samples.

## 1. Introduction

Plant populations are not arbitrary assemblages of genotypes but are structured in space and time [1]. The major evolutionary forces -- such as gene flow, genetic drift and different selection regimes -- are responsible for the observed genetic structure of plant species and populations [2]. Moreover, plant populations can also be profoundly affected by human activities, such as habitat modification, domestication, direct extraction and introduction of invasive species [3]. Determining the relative importance of these forces may be extremely complicated, as several factors may be at work simultaneously [1]. Nevertheless, an understanding of the population genetic structure of a species can yield critical information for conservation, management and general understanding of plant evolution [1,4].

Migration among plant populations is affected by life-history traits (e.g., mating system), pollinators and seed dispersal mechanisms, and by the continuity of geographical distribution [5-7]. Some pollinators, like nectar feeding bats, can travel long distances, therefore reducing the structuring of genetic variation among pollinated populations [8-10]. Whereas, other pollinators, like bees, usually travel short distances [11,12], which would favor mating between closely spaced individuals [13]. Similarly, with seed dispersers, short-distance vectors (e.g., gravity) move seeds only a few meters, thereby limiting migration, whereas long-distance vectors (e.g., migratory birds or bats) move seeds much further, thereby facilitating migration and reducing genetic structure [10,13,14].

Another important factor affecting plant’s population genetic structure is spatial distribution [1,15]. Plant species can occur in uniform, clumped, or random distributions, or, for instance, along an elevation gradient; and the ranges of species may vary in size from widespread to very narrow and endemic. It is thought that plant species with a large or patchy distribution will exhibit stronger genetic structure than those with a small or continuous distribution, because it is more difficult to move pollen and seeds across larger spatial distances [16,17]. Moreover, species distributed across wide heterogenous environment may be exposed to differential selection regimes or to local adaptation [18]. In this case, the successful occupation of a wide area may be explained by the occurrence of many specialized genotypes, rather than by the existence of a single universal genotype. Therefore, divergent selection among these populations inhabiting ecologically different environments would create a barrier to gene flow, thereby promoting genetic differentiation [19,20].

Genetic structure in plants is also affected by human use and cultivation [21,22]. Usually, the most serious impact on the effective population size (*N*_*e*_) and connectivity among plant populations comes from direct extraction and lumbering [21,23]. On the other hand, many plant species that have been historically cultivated may be located outside their natural range, and genetic isolation from source populations may lead to increased genetic structuring. Cultivation of a limited number of individuals may cause a founder effect, resulting in genetic drift, lower genetic diversity and increased differentiation [24]. Artificial selection for desirable traits in the process of domestication may also decrease genetic variation and further increase genetic structure [25]. Otherwise, cultivation may reduce genetic structure if genotypes are obtained from multiple source populations, or if gene flow occurs between cultivated and local wild populations [26,27]. Given the above, determining and explaining population genetic structure of plant species, although very important, may be challenging.

In this regard, species of *Agave* L. (Asparagaceae) may be an excellent model to investigate the relative importance of natural and/or anthropogenetic causes that affect population structure and diversity in plants. *Agave* is a species rich genus of perennial monocots found primarily in the arid regions of Mexico and the south-western United States [28,29]. *Agaves* are keystone species that disproportionately affect their ecological community relative to their abundance by producing a large inflorescence with copious amounts of nectar and pollen [30-32]. On the other hand, *agaves* have a long history of interaction with humans and they have been utilized historically by native populations in North America as a source of fiber, food, beverages and medicine [31,33]. The spirits production from *Agave* has a long history, and nowadays it has become the primary use of agave [29,31]. Over fifty species of *Agave* L. are used for beverages production [29,34]; yet, the majority of the utilized species are not cultivated but rather extracted from natural populations [29,35]. Nevertheless, a growing number of species are now managed [33] with different levels of intensity, from planting wild seeds in backyards to extensive monocultural plantations of laboratory produced clones with no flowering allowed [36]. In the last decades the demand for tequila like beverages has grown dramatically; for example, since 2014 the production of mezcal has increased by almost 500% [37]. This trend has not only increased a pressure on natural populations, but has also introduced industrial level techniques into plant production and gathering [38].

The increase in mezcal production, and in particular the accelerated extraction of wild individuals, constitutes an important environmental problem [39]. This issue is well exemplified by the tequila expansion, which promoted rapid growth of plantations, massive destruction of natural vegetation, soil erosion, reliance on vegetative propagation, and prevention of cross-pollination and gene flow [38]. As a consequence, the genetic diversity of *A. tequilana* was drastically reduced [40, 41], which led to an increased vulnerability to pathogens [42]. The available studies on other agave species indicate that there may be a considerable reduction in genetic diversity in cultivated individuals [41,43,44]. On the other hand, several studies have reported that owing to the incipient stage of domestication and recent formation of crop areas, many agave species still maintain relatively high genetic diversity. However, there is already a trend towards a decrease in genetic variation and increase in differentiation [41,45-47]. Apparently, the reduction of genetic diversity and increased differentiation are related to the intensity and time under management [48]. Given the above, it is of the utmost importance to apply state-of-the-art methodologies to evaluate natural and anthropogenic factors that may affect population structure and diversity of wild and cultivated agave species, and therefore, avoid the loss of genetic variation and evolutionary potential.

In this study we characterized the patterns of genomic diversity at varying geographic scales, and investigated the relative importance of natural forces and human activities in shaping patterns of genetic structure of *A. angustifolia* var. *pacifica* an agave used for bacanora (an alcoholic spirit similar to tequila and mezcal) production *in the Sonora state* in northwestern Mexico. We analyzed 96 individuals from wild and cultivated sites of *A. angustifolia* var. *pacifica*, genotyping them with over 11,000 SNPs markers (Table S1). Our aims were to (1) determine whether and how geographic distribution, life-history traits (e.g., mating system, dispersal mechanisms, longevity) and human activities have affected patterns of genetic structure and diversity in this species; and (2) determine the extent to which we can detect genetic signatures of intensifying human management on the wild and cultivated populations. We hypothesized that due the fact that *A. angustifolia* pollinators may potentially move genetic material across long distances, a shallow population structure among sites and considerable genetic diversity would be observed in wild plants. We also expect that due to the patchier geographical distribution, history of cultivation and possible genetic drift, individuals under management would show less genetic diversity and stronger genetic structure relative to the wild populations. Such an approach has the potential to aid conservation and management strategies because it can identify at-risk, low-diversity wild and cultivated sites that would benefit from restored gene flow within a broader geographic region.

## 2. Results

The GBS on 96 *A. angustifolia* samples resulted in a total of 407,485,082 PE raw reads, with an average of 3,953,325 reads per sample (range 2,006,790 - 5,647,966). After filtering and adapters removal, the final data set resulted in an average of 2,630,790 high quality reads per sample (range 1,114,581 - 3,785,957). From this data, 638,704 variants were called using the *de novo* pipeline. After filtering, our final data set consisted of 95 *A. angustifolia* individuals and 11,619 SNPs with 2.8% of the missing data. Average sequencing depth for each individual and missingness on per site and per individual level are present in the Table S2.

### 2.1. Population structure

Overall, populations of *A. angustifolia* showed shallow genetic differentiation, with the exception of some cultivated sites and one wild site (Figures 1, S1 and S2,). First of all, although different from zero, genetic differentiation between wild and cultivated plants was low. For example, the averaged paired *F*_ST_ was 0.005 (CI at 95 %, 0.0046-0.0058). Similar results were found using Nei’s *D* index, as cultivated plants had negligible differentiation from their wild counterparts (0.006). We further carried out differentiation analysis at site level for cultivated and wild individuals and only for the wild sites (Figures S1and S2). For the complete data set, we found that one wild and three cultivated sites presented comparatively high differentiation from the rest of samples (*F*_ST,_ ranged from 0.1 to 0.57). One of these sites corresponded to a plantation that had *in vitro* propagated plants (MocC3; personal communication with owner; [36]), another corresponded to samples from an intensively managed monocultured plantation and the last corresponded to a back-yard plantation from a small village. Only one of the wild sites (CamW) was highly differentiated (*F*_ST_ > 0.35; Figures S1 and S2) from the rest of the wild samples. After excluding this site (CamW), the differentiation among wild sites was moderate (*F*_ST_ = 0.076, ranging from - 0.01 to 0.22; Figure S2). We also found considerable differentiation between coastal sites (elevation lower than 10 m) and the rest of the samples (*F*_ST_ ranged from 0.09 to 0.22). When these coastal sites were excluded, the differentiation was low and homogenously distributed, ranging from *F*_ST_ -0.01 to 0.01 (Figure S2).

**Figure 1.**
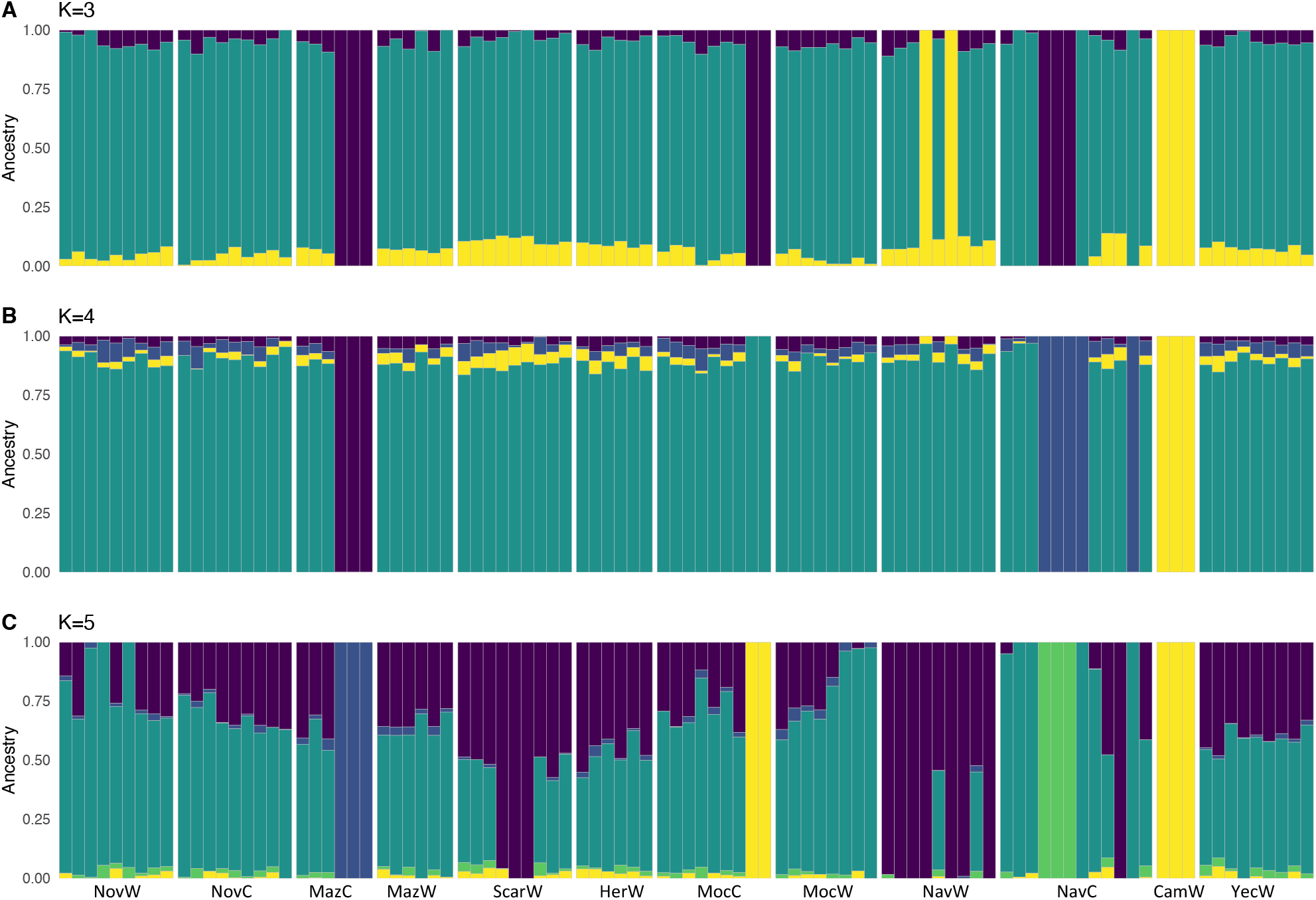
Plotted are modes with clustering solutions obtained with 95 wild and cultivated individuals of *Agave angustifolia* var. *pacifica* from the state of Sonora, Mexico. Values of *K from 3 to 5* are shown. In each plot **(A-C)**, each cluster is represented by a different color, and each individual is represented by a vertical line divided into *K* colored segments with heights proportional to genotype memberships in the clusters. Thin white lines separate individuals from different regions and management types as coded in table S1 (e.g., NovW corresponds to Novillo region and wild samples, whereas MazC corresponds to Mazatan region and cultivated samples).

To investigate the fine scale population structure between and within wild and cultivated *A. angustifolia* we used an ADMIXTURE analysis. When all samples (cultivated and wild) were used, the cross-validation error estimates showed that model fit was optimized at *K* = 4 (Figure S3). Nevertheless, to better understand the genetic structure within the samples, we plotted the results from *K* = 3 to *K* = 5 (Figure 1).

In general, ADMIXTURE analyses found shallow genetic structure within *A. angustifolia var. pacifica, with only one wild site and several cultivated individuals being differentiated*. Based on the complete data set, we were able to group individuals into the following clusters (corresponding to *K*=5; Figure 1C): (i) wild individuals from one site (CamW) plus two cultivated individuals from Moctezuma region (in yellow); (ii) three individuals from a back-yard plantation of the Mazatan region (in dark blue); (iii) three individuals from an intensively managed plantation near Navojoa (in green); (iv) ten wild individuals from the coastal area near San Carlos and Navojoa (in purple); and (v) all the rest of the cultivated and wild samples. Some individuals from the last group (v) also presented mixed ancestry with samples from the coastal group (iv) (Figures 1A, B, C).

A PCA showed patterns consistent with those described by the pairwise *F*_ST_ comparisons and ADMIXTURE clustering (Figure 2).

**Figure 2.**
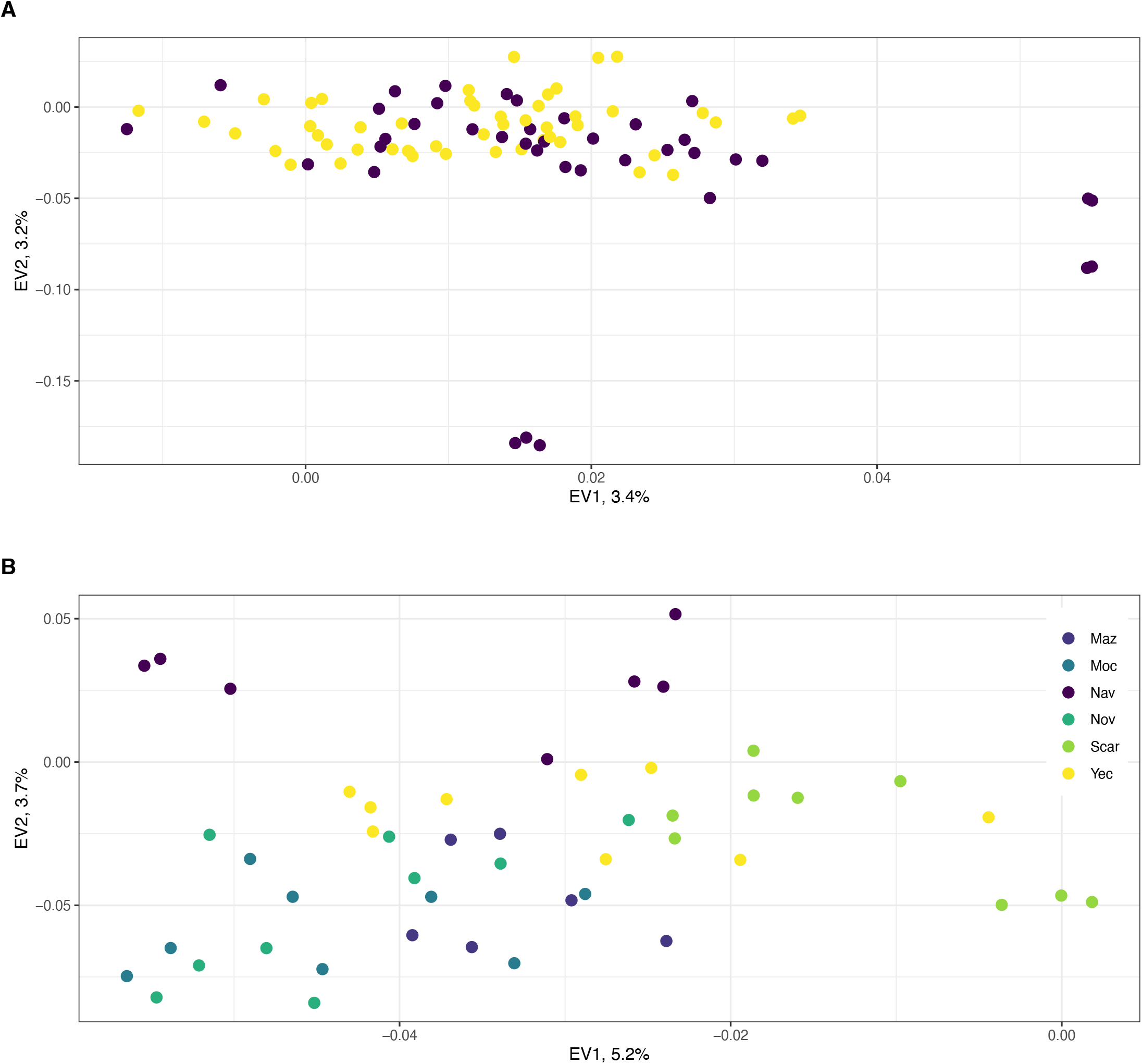
Relationships among **(A)** *A. angustifolia* var. *pacifica* from the state of Sonora, Mexico, wild (yellow) and cultivated (purple) samples as represented by principal component analysis (PCA) using 11,619 genome-wide SNPs and excluding 6 outlier samples. Panel **(B)** PCA representing relationships among wild samples of *A. angustifolia* var. *pacifica*. Colors corresponds to the major geographic regions detailed in table S1.

After removing outlier samples (three wild individuals of CamW population and tree samples from the back-yard cultivated plants from Mazatan region, Figure S4), and except for a handful of cultivated samples, all individuals clumped into one cloud, with no apparent differentiation between management type or site (Figure 2A). The cultivated samples that were separated from the main cluster were: (i) *in vitro* plants and (ii) plants from one of the intensively managed plantations near Navojoa city. When we focused only on the wild samples (Figure 2B), in the first eigenvector (which explained 5.2 % of the variance) most individuals formed a cloud, with some separated individuals from the coastal site of San Carlos region. The second eigenvector (3.7%) further separated coastal samples from the southernmost sites near Navojoa from the rest of the samples.

### 2.2. Spatial genetics

For the spatial analysis, we focused only on the wild samples, also excluding one sampling site (CamW) that presented higher differentiation. The cross-validation criterion performed with *TESS3R* did not exhibit a minimum value or a plateau (Figure S5), but there was a steady increase in statistical support at higher *K* values. However, after *K*=5 biologically meaningful structure was lost. Therefore, we plotted the results of different values of *K* (from *K*=2 to *K*=4; Figure 3).

**Figure 3.**
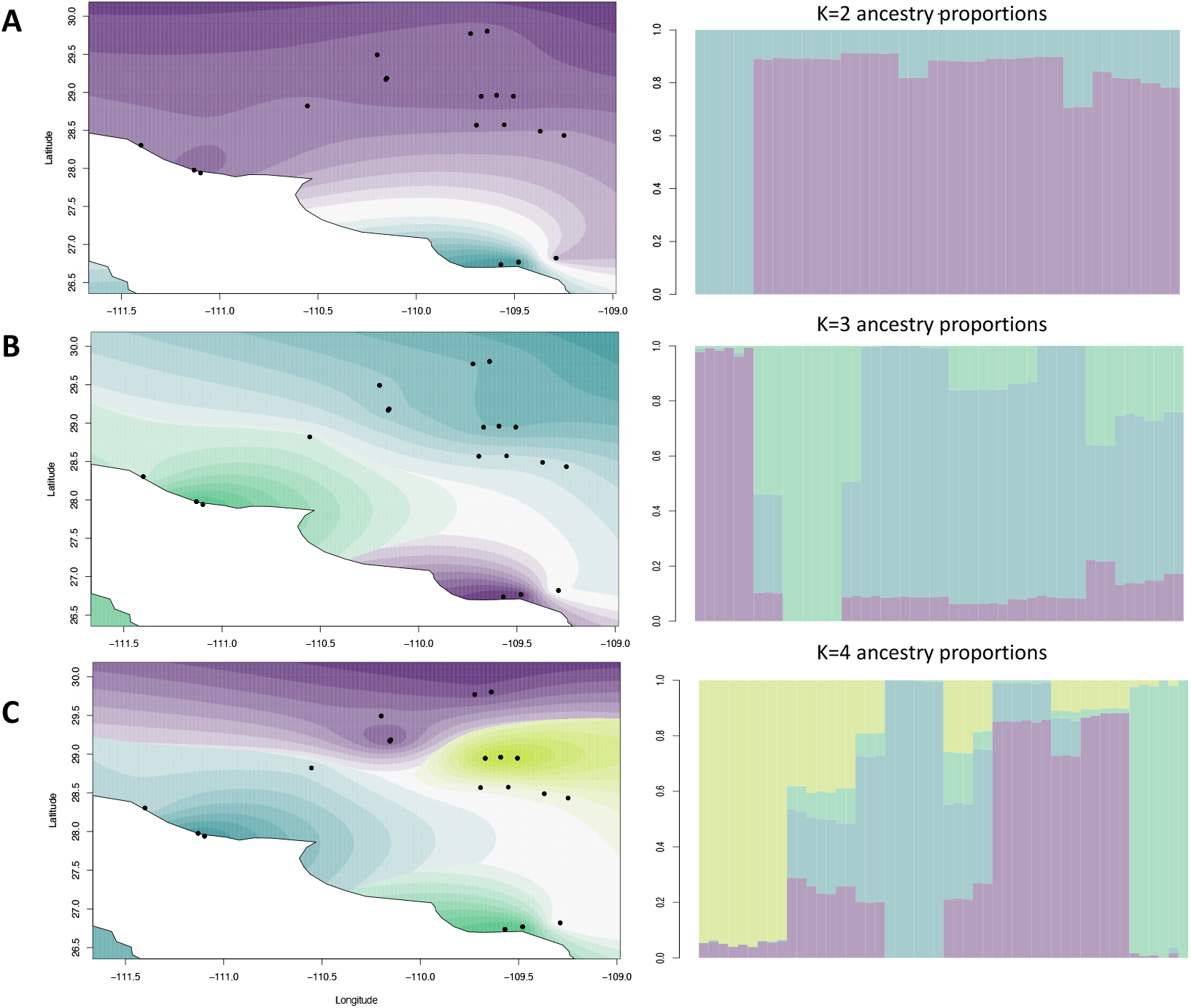
Interpolated ancestry proportions from TESS3R, demonstrating the geographic distribution of biologically meaningful genetic clusters (*K*) from 2 to 4 in *Agave angustifolia* var. *pacifica* from the state of Sonora, Mexico. Locations on each map are colored by the resulted dominant ancestry cluster, with transparency reflecting the percent ancestry of that cluster, with the largest value assigned as opaque.

At *K*=2 (Figure 3A) we found support for differentiation of two coastal southernmost sites. Further partitioning at *K*=3 added the San Carlos region coastal samples as a separate group (Figure 3B). At *K*=4, we observed differentiation of the central sites collected between Novillo and Bacanora villages, that was not observed with any other analysis (Figure 3C). Spatial PCA analysis revealed highly significant global (*p = 0.001*) but non-significant (*p = 0.89*) local spatial structures (Figure S6), indicating signatures of among sites separations. A plot of lagged scores from the first two principal component suggested that the global structure was linked to the elevation and differences of the coastal environment (Figures S7), corroborating with the *TESS3R* results and pairwise *F*_ST_. Taking together, we interpret that our sampling area has at least three genetic clusters: (i) samples of two coastal sites from low elevation near Navojoa city; (ii) group including three coastal sites of low elevation near San Carlos; and (iii) an inland group comprised by the rest of the sites.

Genetic distance (*F*_ST_) between wild sampling sites increased with geographical distance (Mantel test = 0.41, *p*= 0.007); nevertheless, the relationship was not linear (Figure S8A). Moreover, a plot of *F*_ST_ vs. elevation distance revealed less strong but still significant relationship (Mantel test *r* = 0.19, *p*= 0.01; Figure S8B).

### 2.3. Genetic diversity

The overall mean observed heterozygosity (*Ho*) for all samples was 0.22 (SD 0.13). The mean *Ho* among 53 wild agave individuals was lower (0.22, SD 0.12) than the expected heterozygosity (*He* = 0.25, SD 0.13), indicating a deficit of heterozygous individuals. Almost identical results were obtained for the cultivated samples (*Ho*=0.22, SD 0.13 and *He* = 0.25, SD 0.13). The difference between expected and observed heterozygosity was significant, even after Bonferroni correction (Bartlett test all samples K-squared = 84.965, *p-value* < 2.2e-16; only wild samples K-squared = 36.168, *p-value* = 1.81e-09). Individual based multilocus heterozygosity (MLH) was similar to the *Ho* for both, wild and cultivated plants (Table S3). In addition, two frequency-weighted measures of individual heterozygosity – standardized multilocus heterozygosity (sMLH) and internal relatedness (IR) – showed no difference between wild and cultivated individuals (Table S3 and Figure S9).

We found that wild and cultivated agave plants are moderately inbred according to the Fhat3 inbreeding index, with similar *f*, averaging 0.13 (SD 0.01) for both wild and cultivated agaves. In wild individuals *f* ranged from almost zero to as high as 0.37 (Table S3 and Figure S9); while in cultivated plants, inbreeding ranged from 0.09 to 0.24 (Table S3 and Figure S9). Similar results were obtained using Wright’s *Fis* (Table S3).

When samples were partitioned at the population/region level and management type, we found similar pattern of homogeneous distribution of heterozygosity and inbreeding (Table S4). The lowest inbreeding, 0.11 was found in wild populations of Navojoa and Mazatan, whereas the highest estimates (0.15) were found in two wild populations and one of cultivated plants (Table S4). Nevertheless, after FDR correction, none of the comparisons was significant. Further partition of wild samples was made based on the spatial clustering of the samples recovered with *TESS3R* (Table 1). We found significant (*p=0.02*) difference in MLH and *F*_*is*_ between inland and coastal sites; with coastal sites presenting lower diversity and higher inbreeding (Table 1 and Figure S10).

**Table 1.** Diversity estimates and standard deviation (in parenthesis) as estimated for wild and cultivated *Agave angustifolia* var. *pacifica* in the state of Sonora, Mexico. Wild populations were determined based on spatial analysis in TESS3R. One highly divergent wild site (Cam) is excluded from this table. N - number of individuals; sMLH - standardized multilocus heterozygosity; MLH - multilocus heterozygosity; IR - internal relatedness; *Fis* - Wright’s inbreeding index; Fhat3 - inbreeding index.

| Diversity index | Wild Populations |  |  | Full dataset wild | Cultivated |
| --- | --- | --- | --- | --- | --- |
|  | South Coastal | Central Coastal | Inland |  |  |
| N | 6 | 9 | 35 | 50 | 42 |
| sMLH | 1.00 (0.063) | 0.98 (0.011) | 1.00 (0.018) | 0.99 (0.014) | 1.00 (0.03) |
| MLH | 0.22 (0.013) | 0.22 (0.002) | 0.22 (0.004) | 0.22 (0.003) | 0.22 (0.002) |
| IR | 0.05 (0.048) | 0.06 (0.007) | 0.06 (0.014) | 0.05 (0.011) | 0.05 (0.02) |
| Fis | 0.13 (0.054) | 0.14 (0.009) | 0.13 (0.016) | 0.13 (0.012) | 0.13 (0.03) |
| Fhat3 | 0.11 (0.047) | 0.13 (0.001) | 0.13 (0.017) | 0.13 (0.012) | 0.13 (0.02) |

The mean relatedness was of 0.007 (CI 0.002) (Figure 4), indicating very low relationship among most sampled individuals. Nevertheless, we did find high (over 0.5) levels of relationship between some of the comparisons (12 pairs of individuals). These presumable clones were represented by plants sampled at intensively managed (NavC) and the *in vitro* propagated (MocC) populations.

**Figure 4.**
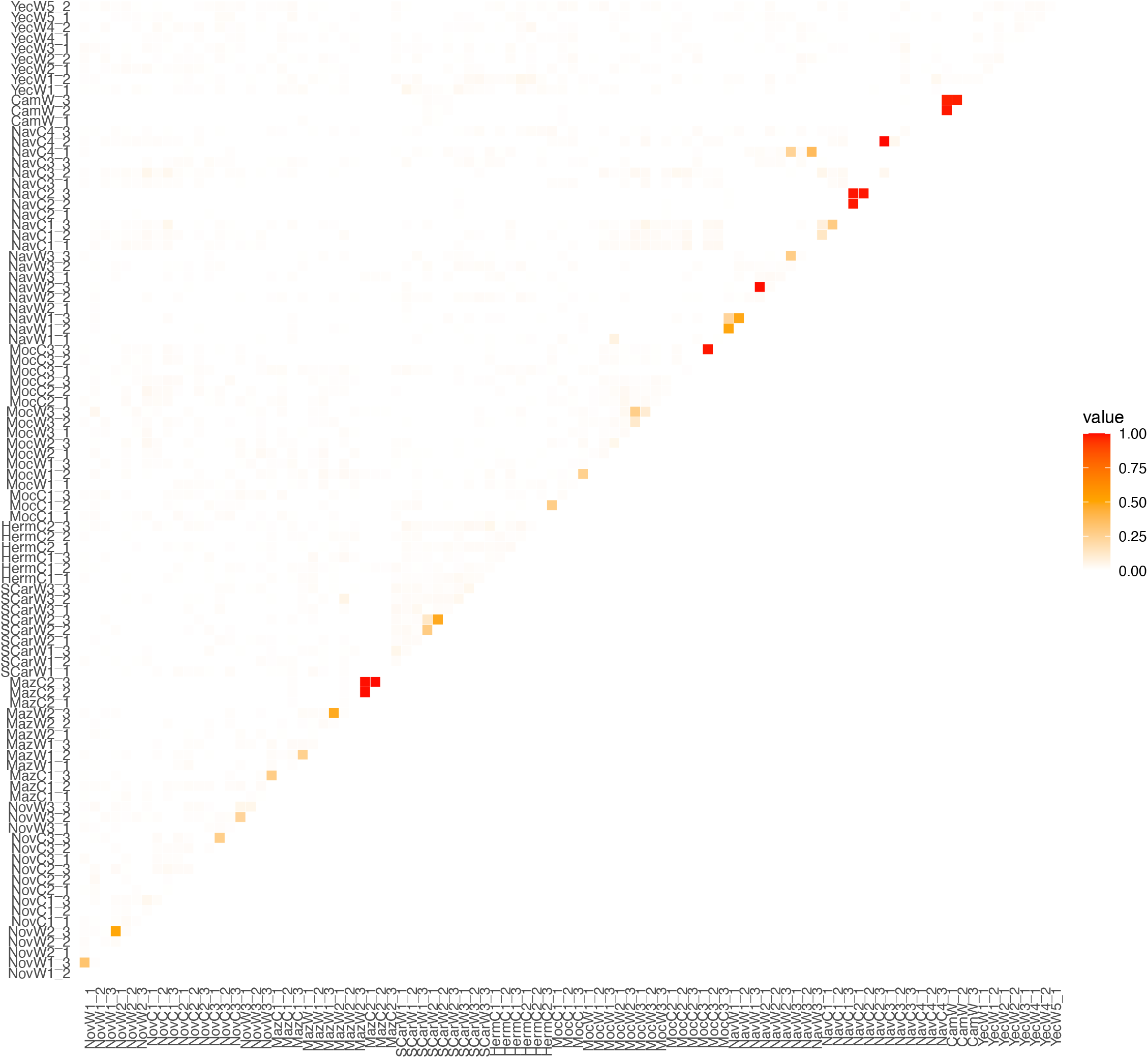
The overview diagram of relatedness coefficient as estimated with TrioML methodology among 95 agave individuals *Agave angustifolia* var. *pacifica* from the state of Sonora, Mexico. The degree of relatedness is represented by colors, from white that indicates no relationship, to red representing a clonal relationship. Samples are coded according to the Table S1.

When the number of private alleles was compared between managed and wild plants, we found that wild individuals harbor a higher number of alleles that were not shared with cultivated plants (i.e., “private” alleles), 299 vs. 12.

## 3. Discussion

We analyzed patterns of genetic diversity and population structure of wild and cultivated *A. angustifolia* var. *pacifica* from northwestern Mexico. *Agave angustifolia* is a species with wide distribution and a long history of human use, that recently has escalated up to intensive management and heavy pressure on wild populations (Molina Freaner pers. observ.). Nevertheless, probably due to a combination of factors, including aspects of the species biology (e.g., long life cycle, monocarpic reproduction), pollination biology (involving bats and other highly mobile animals), the incipient cultivation history and particularities of bacanora production (a spirit less known in the market than tequila and mezcal, lower production and high reliance on wild individuals); we found low genetic structure, and homogeneously distributed levels of diversity in both wild and cultivated individuals. As for the wild samples, we found low but detectable spatial population genomic structure, apparently driven by the ecological characteristics of the habitat.

### 3.1. Population structure in wild and cultivated agaves

One of the aims of this study was to understand how naturals forces, life-history traits and cultivation practices may have affected patterns of genetic structure of *A. angustifolia*. We found that wild populations of *A. angustifolia*, with a relatively continuous distribution in dry, mid-elevation areas throughout northwestern Mexico [49], do not present strong population genetic structure; which is not surprising, considering that most agaves have low genetic differentiation [43].

Overall, genetic structure is created by limited migration among populations, genetic drift and different selection regimes occurring across populations [2]. In *A. angustifolia* var. *pacifica*, shallow genetic structure is probably the result of high genetic connectivity, which homogenizes populations and reduces the effects of genetic drift and selection. Species of nectar-feeding bats in the genus *Leptonycteris* have been proposed as important pollinators of agaves in general, and of *A. angustifolia* in particular [50, 51]. Consistently, agave species pollinated by bats present similarly low population genetic structure [43,44,46]. Interestingly, these bat species themselves have a wide distribution and general panmixia with low genetic differentiation among colonies [51-53].

Another important life-history trait that may contribute to the observed lack of differentiation is the possibility of clonal reproduction and, as a consequence, enhanced longevity of genets [31,49,54]. In the harsh desert environment temporal gaps between years with successful sexual recruitment can be highly variable. Therefore, plants can benefit from clonal reproduction, as clonality can enhance genet longevity, compensate for the partial loss of genets due to disturbance, reduce effective population size fluctuations and, thus, decrease the effects of genetic drift [31,49,55]. Moreover, high longevity of clones may imply that even if the establishment of seeds dispersed over long distances is rare, and individuals arising from seeds dispersed over long distances are outnumbered in a local population, the likelihood of detection and sexual reproduction of such individuals are increased, because they persist for longer periods of time [56].

Although we did not detect population genetic structure using traditional methods, we did find interesting spatial clustering of the samples. Thus, we detected that low elevation coastal sites (with an elevation of less than 50 meters) are differentiated from the inland genetic cluster. Moreover, those sites were also differentiated from each other, the fact that also may explain non-linearity in Mantel tests. Coastal regions have unique environmental characteristics, such as high salinity, greater availability of atmospheric moisture, wind strength and tidal influence, therefore constituting distinct landscapes with unique abiotic and biotic composition [57,58]. We therefore hypothesize that different selection regimes occurring across populations in the coastal environment may be responsible for the observed differentiation. Further studies based on loci located in coding regions or focusing on transcriptome analysis would be crucial in determining genomic regions involved in promoting this low, but significant differentiation [59].

The genetic differentiation between wild and cultivated agaves has been the focus of extensive research [29,43], with variable results. For example, some intensively cultivated varieties and species, such as *A. tequilana*, presents considerable genetic differentiation from its wild relatives and among cultivars [48,60]. Basic explanation for the observed structure includes reproductive isolation through asexual reproduction and high genetic drift. On the other hand, insignificant divergence related to management was observed in other agave species [43,47]. We found that, with the exception of few managed sites, there is no genetic differentiation between wild and cultivated agaves in northwestern Mexico, in other words, the cultivated plants are basically a subset of the wild populations. Cultivation history of *A. angustifolia* var. *pacifica* is relatively recent and therefore, there may have been little time for differentiation.

Our findings expand the knowledge of genetic differentiation and diversification processes in agaves across coastal and inland areas of northwestern Mexico. Our results are concordant with previous suggestion [1,54] that multiple processes are likely influencing genetic structure in natural plant populations. We suggest that pollinator-mediated gene flow and clonal reproduction may serve as homogenization forces, whereas different selection regimes may be responsible in promoting local adaptation and intraspecific differentiation. Our results suggest that a complex mixture of features related to life history traits, environment and management shape contemporary genetic differentiation patterns of *A. angustifolia* var. *pacifica*.

### 3.2. Genetic diversity

The most widespread and concerning consequence of plant management and domestication is the erosion of genetic diversity [61,62]. During the initial phases of domestication and due to the effects of genetic bottleneck, genetic variation is usually reduced through decreasing effective population size and increasing inbreeding [25]. Then, additional loss of diversity is observed due to artificial selection through the removal of allelic variants of genes underlying traits that are undesirable for cultivation [63,64]. To date, genetic erosion and reduction in diversity of gene expression have been observed in many intensively cultivated plant species [65,66].

The case of agave is probably more complicated, with some cultivated varieties (e.g., *A. tequilana, A. salmiana* and *A. americana*) presenting low levels of diversity [40,48,67,68], whereas in other species levels of genetic variability are indistinguishable or even higher in cultivated individuals in comparison to their wild counterparts [43,47]. Here, we found that cultivated individuals of *A. angustifolia* in northwestern Mexico have not lost genomic diversity, nor did they present increased inbreeding in comparison to their wild counterparts. These findings are consistent with other studies reporting that traditional management of agave that involves gathering of wild specimens and seeds, instead of clonal propagation, allows the persistence of high genetic diversity in cultivated agaves [41,46,47]. On the other hand, in many traditional agave plantations, the diversity can be even greater than in wild individuals, due to the different origins of the germplasm introduced into cultivation and to non-strict selection [41,69]. This was also true in our study, since we found considerable genetic divergence (*F*_ST_ >0.3) of some cultivated sites and individuals, particularly at some of the intensively managed plantations. Unfortunately, we were not able to determine the original geographic source of these samples, as probably they were moved from locations outside the sampling area. We suggest that future studies should focus on analysis of *A. angustifolia* throughout its whole distribution area and to establish a standard set of markers that would be useful to trace the origin of the cultivated individuals and to compare their diversity levels.

The incipient domestication in agaves may be targeting particular genomic regions [64,70], and therefore, the putatively neutral markers used here may be not efficient enough to identify small differences between wild and cultivated plants. Future studies may consider markers located at coding regions or transcriptome analysis [71,72]. Additionally, recently developed pangenome analysis may offer an exciting opportunities and high precision in identifying specific regions and genes that are being lost/gained during the process of domestication and management [73,74].

Importantly, although we found that heterozygosity and inbreeding estimates were similar between wild and cultivated individuals, wild samples harbored a larger number of unique (private) alleles (299 vs. 12), which represents an important gene pool of standing genetic variation not found in the cultivated plants used for bacanora production. This genetic resource could be relevant, for example, in future crop improvement programs. These differences indicate that the cultivated plants do not include the total genomic diversity of *A. angustifolia*, since they are just a small sample of the diversity found in wild agaves.

### 3.3. Inbreeding

In contrast to the general expectation that the lowest levels of inbreeding in plants would occur in outcrossing species, a significant excess of homozygotes and moderate levels of inbreeding (average *f =* 0.13 for both wild and cultivated plants) were observed, and these occurred consistently among sites. Interestingly, in the *Agave* genus, relatively high levels of inbreeding (*F*_IS_) have also been reported in other studies [43]. For example, considerable inbreeding was reported in wild and cultivated samples of *A. maximiliana* (*F*_IS_ range from 0.002 to 0.32; [47]), in *A. americana, A. salmiana* and *A. mapisaga* (*F*_IS_ ranged from 0.19 to 0.72 [68]) and in *A. potatorum* (*F*_IS_ ranged 0.21-0.32; [75]).

Explaining moderate levels of inbreeding coupled with considerable polymorphism, that have been found in many plants, has not been an easy task [76]. One of the interesting hypothesis states that individual reproductive variation and temporal variation in flowering among individuals, coupled with the long life of plants, may split population into sublines, with occasional mixing of sublines. Within sublines, inbreeding and genetic drift are increased, but genetic drift of the entire population is decreased. Therefore, the sublines would maintain both high levels of polymorphism and explain the homozygote excess [77]. Moreover, Trame et al. [78] estimated that inbred progeny of *Agave schottii* apparently does not experience strong negative selection and, as a consequence, inbred individuals continue to make up a substantial proportion of the population. Nevertheless, this hypothesis has not been tested, in *A. angustifolia*, and further research focusing on optimal outcrossing distance and effects of inbreeding depression in this species would be helpful in elucidating how moderate levels of inbreeding and low population structure can be maintained [78,79].

### 3.4. Conservation and management implications

The major concern of the current increase of mezcal and related beverages production is that in the eagerness of satisfying ever-growing demand, there could be an unprecedent negative impact on natural habitat and on wild agave populations [47,80]. Currently, it is difficult to evaluate *A. angustifolia’s* level of genetic diversity in a broader context of Agavoideae species, and it is unclear what levels of diversity and inbreeding should be expected. These challenges result from the general lack of studies that cover the whole distribution range of this or other agave species and because of a myriad of different molecular markers that have been used over the last 50 years [29,43].

Nevertheless, we have discovered an alarming trend in some cultivated samples, particularly collected at the intensively managed plantations.

Although, at each plantation we sampled distant individuals, we found several samples with high relatedness coefficient (over 0.5), indicating clonal propagation. These findings are in contrast with the fact that almost all wild individuals and samples from the traditionally managed plantations were genetically unrelated. Clonal propagation has several advantages for cultivation, for example, it is easier and faster than propagation from seeds, it also ensures that favorable genotypes and mutations are passed on to the next generation [81]. Nevertheless, in a long-term absence of sexual recombination under exclusive clonality, would lead to erosion of genetic diversity and loss of evolutionary potential [82]. Without recombination, the mutational load increases, that ultimately would lead to lower fitness and decrease in agronomic performance [81,83]. We, therefore, argue that if the reliance on vegetative propagation would increase, ultimately it may be detrimental to bacanora production [84].

Based on our findings we argue that there is an urgent need to conduct systematic research focused on (i) delineating distribution limits of agave species used for spirits production; (ii) determining conservation and management units within each species; (iii) evaluating the impact of wild plant extraction by humans on species distribution and numbers; (iv) developing a set of genomic markers that would allow reproducibility and comparison among studies. Moreover, ancient and historical genetic data, such as those from specimens stored in herbarium collections and remains from archaeological sites, can add a temporal dimension to conservation of agaves by providing baseline levels of diversity [85].

## 4. Materials and Methods

### 4.1. Study species

*Agave angustifolia* is a diploid, *apparently self-incompatible [50,86]* species *with high rate of sexual reproduction* and the widest geographic distribution within the genera [31]. *It is pollinated by Leptonycteris bats*, birds and bees [50]. *It* is widely and relatively continuously distributed from Sonora and Tamaulipas in Mexico to Costa Rica [31]. It occupies dry shrublands, tropical dry forests, chaparral and pinyon-oak woodland habitats, and can be found from sea level to an elevation of 2,500 m [31,87]. Because of its wide distribution and unresolved phylogenetic relationships, *A. angustifolia* is sometimes considered to comprise several varieties [69]. In northwestern Mexico, particularly in the state of Sonora, the complex is represented by *A. angustifolia* var. *pacifica*, which is the focus of the present study.

*Agave angustifolia* has a rich ethnobotanical history, and it is among the most important *Agave* spp. utilized in Mexico [31,49]. Ethnobotanical records suggest that humans in northwestern Mexico have utilized *A. angustifolia* [88,89], but the species has no documented history of cultivation for distilled spirits production until arrival of the Spanish settlers. The distribution pattern of *A. angustifolia* in northwestern Mexico is island-like, as it is mostly found in scattered populations upon the rocky slopes of hills and mountains. It is also widely distributed in the coastal thorn forest, where it grows on sandy soils and gravelly terraces [49]. Currently, in Sonora *A. angustifolia* is used for bacanora production [90], and in some areas it is actively cultivated. Cultivation is now often carried out in monocultural extensive areas. In general, cultivated plants are prevented from flowering.

### 4.2. Sample collection

We analyzed 96 adult plants of *A. angustifolia* collected in 34 sites in the state of Sonora, Mexico in October 2021 (Figure 5 and Table S1).

**Figure 5.**
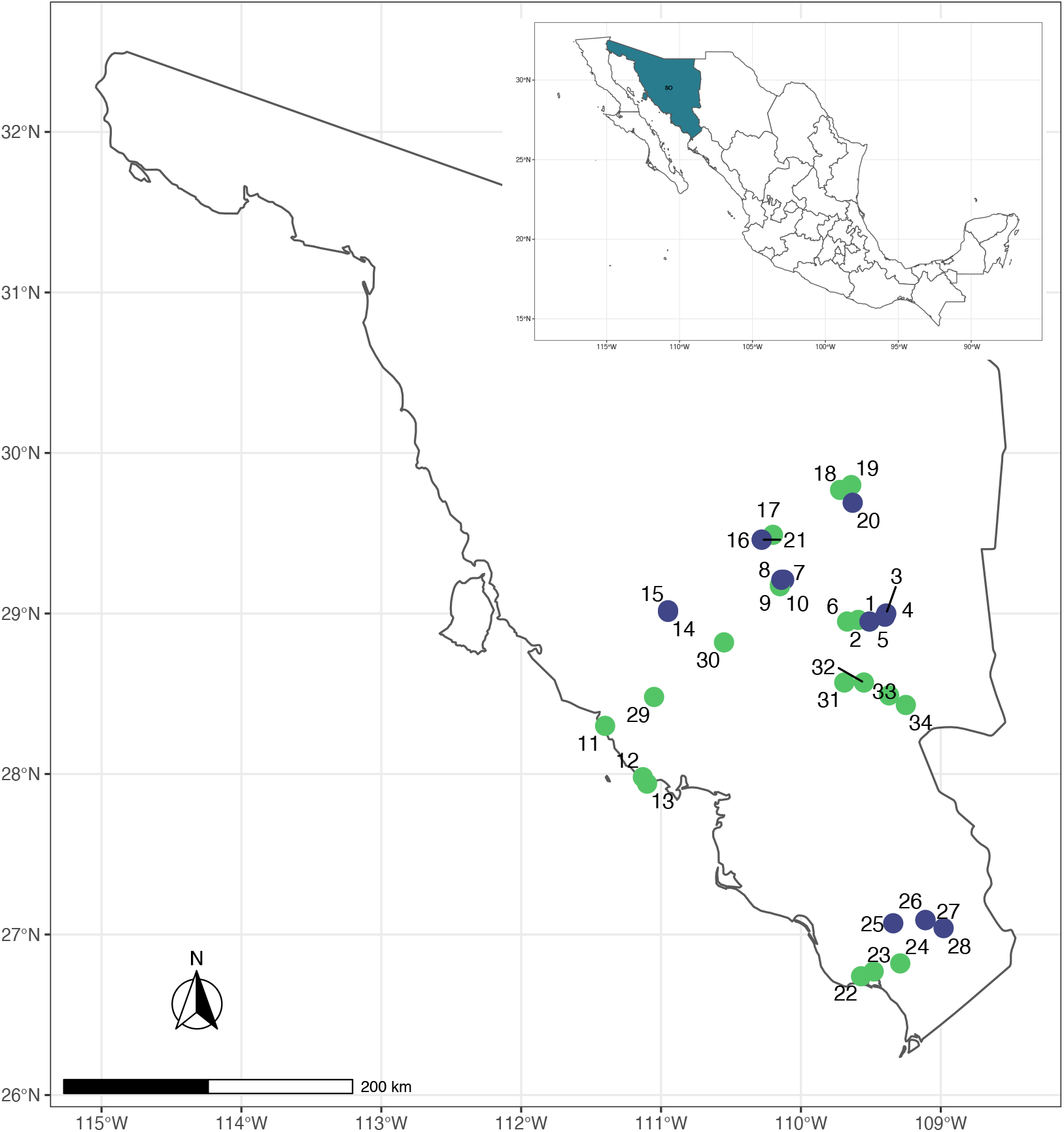
Map showing sampling sites of wild (green) and cultivated (blue) *Agave angustifolia* var. *pacifica* from the state of Sonora, Mexico. The full names of each site and sample sizes are given in Table S1.

From those, 20 sites belonged to wild populations, 11 sites represented Bacanora plantations, and three sites were backyard plantations of *A. angustifolia*. More details on geographic coordinates, number of specimens collected at each site and management type can be found at table S1. Fresh material from the field and cultivated specimens was stored at -20° C until DNA extraction.

### 4.3. DNA extraction, library preparation and sequencing

For DNA extraction, three individuals were chosen randomly from each sampling site to reach a total of 96 plants. Total genomic DNA was extracted from disrupted lyophilized with liquid nitrogen leaf tissue using a modified CTAB protocol [91,92]. Total genomic DNA was checked for degradation using a 1.5% agarose electrophoresis gel. The quantity of the DNA was determined using Qubit 3.0 fluorometer with Qubit dsDNA broad-range kit. Samples standardization and libraries preparation were performed at the University of Wisconsin Biotechnology Center. Each DNA sample was digested using a combination of methylation -sensitive restriction enzymes (PstI/MspI). The choice of enzymes was based on a previous standardization for *Agave salmiana* and *Agave lechuguilla*. A unique barcode was ligated to each sample, after that all samples were combined and sequenced at the University of Wisconsin Biotechnology Center using NovaSeq 2×150.

### 4.4. Bioinformatics analysis

The raw data were filtered by removing adapters and low-quality bases using TRIMMOMATIC [93]. For the initial filtering the following parameters were used: LEADING:25, TRAILING:25, SLIDINGWINDOW:4:20, MINLEN:60. After first quality filtering, reads in FASTQ format were demultiplexed, filtered and RAD loci were *de novo* assembled using ipyrad v. 0.9.79 [94] with parameters recommended for paired-end GBS data (https://ipyrad.readthedocs.io/). As no published genome of any *Agave* species is yet available, we used a *de novo* assembly strategy. We set the level of sequence similarity for clustering at 90%, after comparing the number of recovered single nucleotide polymorphisms (SNPs) using values between 85 – 95% (see Table S5). The final data filtering was performed using VCFtools v.0.1.15 [95]. Only loci with a mean minimum depth (across individuals) of over 14 and maximum 2 alleles with no InDels were kept. Additionally, we set a minor allele frequency at 0.05, to reduce the possibility of removing true rare alleles that are important in elucidating fine-scale structure [96]. We excluded sites on the basis of the proportion of missing data, keeping sites with no more than 10% missing data (--max-missing 0.9). One individual was removed from the dataset due to high percentage of the missing data (i.e., greater than 50%), which was likely caused by poor DNA quality. We then filtered out the variants that significantly deviated from HWE (p ≤ 0.05 after Bonferonni correction). Finally, to ameliorate the confounding effects of linkage disequilibrium (LD), we eliminated markers within the specified distance from one another, using --thin argument (--thin 250) as implemented in VCFtools. The final VCF file was then produced for all downstream analyses.

### 4.5. Patterns of genetic structure in wild and cultivated individuals

We used several complementary approaches to test for patterns of genetic structure between wild and cultivated agave plants and among sites. We estimated genetic distance between wild and cultivated samples, and among all pairs of sites using *StAMPP* package [97]. Clustering among samples without *a priori* grouping was inferred using the method implemented in ADMIXTURE v.1.23 [98,99]. Admixture analysis was run using 10,000 bootstraps, the number of clusters was set from 1 to 12 (*K*), with ten replicates for each *K* value. The support for different values of *K* was assessed according to the likelihood distribution (i.e., lowest cross-validation error), as well as by visual inspection of the co-ancestry values for each individual. We also used the R package *SNPRelate* to perform a principal component analysis (PCA) of *A. angustifolia* individuals based on the genetic covariance matrix calculated from the genotypes [100]. We ran two separate PCAs for *A. angustifolia* samples: (i) including both wild and cultivated plants, and (ii) including only wild individuals.

### 4.6. Spatial genetics

To explore the role of geography in population genetic structure of *A. angustifolia*, we used spatially explicit clustering program *TESS3R* [101]. *TESS3R* determines genetic variation in natural populations considering simultaneously genetic and geographic data. We tested *K* = 1–10 possible genetic groups with 20 replicates of each *K*. We kept the most supported model (i.e., “best” based on cross-entropy scores) within each of the 20 replicates. Map locations were colored by the resulted dominant ancestry cluster, with transparency reflecting the percent ancestry of that cluster, with the largest value assigned as opaque.

While classic PCA is appropriate to detect obvious genetic structure [102], it does not take spatial information into account, and may therefore miss cryptic spatially arranged genetic structure. In order to confirm and visualize the genetic structure detected by spatial clustering in *TESS3R*, we conducted a spatial Analysis of Principal Components (sPCA) [103]. This procedure is a spatially explicit multivariate analysis that allows to focus on part of the genetic variation that is spatially structured by optimizing not only the genetic variance between samples, but also their spatial autocorrelation. sPCA was conducted only for the wild samples using the package *adegenet* [103]. Tests for global and local genetic structure according to the definition given by Thioulouse et al. [104] were made using 1,000 permutations. The visualization of detected structure was performed by plotting the samples according to their geographic coordinates, and coloring them according to their respective scores along the first and second sPCA components.

Finally, we employed a Mantel test using *ade4* package to estimate the correlation between the genetic distance matrix, geographical, and altitudinal distance matrices [105]. The test uses Pearson’s regression coefficient between distance matrices and 10,000 randomizations. The statistics and the test’s *p* value correspond to the proportion of times the randomized regression coefficient is equal or greater than the observed one. We calculated the pairwise genetic distance matrices among sites using *F*_ST_ and pairwise geographical distance matrices using the Geographic Distance Calculator [106].

### 4.7. Genetic diversity

Expected heterozygosity (*H*_*e*_) and observed heterozygosity (*H*_*o*_) were calculated for the full dataset, and separately for each management type using the R packages *adegenet* and *hierfstat* [103,107]. We also calculated multilocus heterozygosity for each individual, which is defined as the total number of heterozygous loci in an individual divided by the number of loci evaluated in the focal individual [108]. Further, two frequency-weighted measures of individual heterozygosity – standardized multilocus heterozygosity (sMLH) and internal relatedness (IR) were quantified using R packages *Rhh* and *inbreedR* [108,109]. We used plink 1.9 [110] to calculate the inbreeding index Fhat3 [111,112] and Wright’s *F*_IS_ statistics. Following Marshall et al. [113]; we designated inbreeding coefficients (*f*) of zero as ‘none’, below 0.125 as ‘low’, 0.125 ≥ *f* <0.25 as ‘moderate’, and *f* ≥ 0.25 as ‘high’. Wilcox tests were then used to test for significant differences in the diversity indices between management types, sites and populations identified with spatial analysis in *TESS3R* using R package *stats* [114]. We also estimated relatedness coefficient between each pair of individuals using TrioML methodology as implemented in COANCESTRY v. 1.0.1.10 [115,116]. For relatedness analysis, we used 100 bootstraps and accounted for inbreeding using 100 reference individuals. Finally, we estimated the number of private alleles in wild and in cultivated plants using *poppr* package [117].

## 5. Conclusions

The current study aimed to dissect the genetic architecture of bacanora agave (*A. angunstifolia* var. *pacifica)* from northwestern Mexico using SNPs markers. We focused on wild and cultivated sites composed from individuals of the morphologically predominant morphotypes. Nevertheless, even this criterion did not help us from sampling morphologically similar but genetically different plants at one wild site (CamW) and several cultivated sites. Unfortunately, due to the considerable gap in agaves phylogenetic and population genetic studies, we could not trace the geographic original source on these genotypes. Besides this outlier cultivated samples, we did not detect genetic differentiation between wild and cultivated individuals, neither we found loss of genetic diversity in plants under management, suggesting that seeds of wild individuals are readily used for cultivation. Interestingly, the data analysis of wild samples revealed a partition of the samples into three groups—two coastal groups and one inland cluster. Nevertheless, being aware of our experimental limitations, in this moment we are unable to distinguish between local adaptation and neutral divergence; we therefore may only hypothesize reasons for such a separation.

## Supplementary Materials

Table S1: Sites where individuals of *Agave angustifolia* var. *pacifica* from the state of Sonora were collected, including information on geographical region, site abbreviation, coordinates, number of individuals used for genomic analyses (*N*) and sample source (wild, cultivated, backyard plantation). Note: values of *N* do not include one individual removed during the genomic filtering step. All samples were collected in 2021; Table S2: Quality information for the filtered SNPs used in the present study. The data shown is for 95 individuals of *Agave angustifolia* var. *pacifica* from the state of Sonora, Mexico, and 11, 619 SNPs; Table S3: Descriptive genomic diversity estimates and corresponding confidence intervals in parenthesis as determined for each management type. N - number of samples; MLH – multilocus heterozygosity; sMLH – standardized multilocus heterozygosity; *F*_IS_ -Wright’s inbreeding index; Fhat3 - inbreeding index; IR - internal relatedness; Table S4: Descriptive genomic diversity statistics and corresponding confidence intervals in parenthesis for *Agave angustifolia* var. *pacifica* from the state of Sonora, Mexico based on population and management type. N - number of samples; MLH – multilocus heterozygosity; Fis -Wright’s inbreeding index; Fhat3 - inbreeding index; IR - internal relatedness. Region/population column is coded accordingly to Table S1; Table S5: Number of loci and raw SNPs recovered for the datasets under different ipyrad clustering parameters. Figure S1: Pairwise *F*_ST_ differences among all the sampling sites of wild and cultivated *Agave angustifolia* var. *pacifica* from the state of Sonora, Mexico. Colors represent *F*_ST_ values from the lowest of -0.07 in green to the highest of 0.57 in orange. Sampling sites are coded accordingly to Table S1; Figure S2: Pairwise *F*_ST_ differences among all the sampling sites of wild *Agave angustifolia* var. *pacifica* from the state of Sonora, Mexico. Colors represent *F*_ST_ values from the lowest of -0.01 in green to the highest of 0.41 in orange. Sampling sites are coded accordingly to Table S1; Figure S3: Plot of ADMIXTURE cross validation error from *K*=1 through *K*=12, as based on all samples (cultivated and wild); Figure S4: Relationships among *Agave angustifolia* var. *pacifica* individuals from the state of Sonora, Mexico, wild and cultivated samples as represented by principal component analysis (PCA) using 11,619 genome-wide SNPs. All 95 samples were included; Figure S5: Plot of TESS3 cross validation error from *K*=1 through *K*=10, based only on wild samples. Twenty runs were performed for each value of *K*; Figure S6: Results of Eigen value test for sPCA analysis. The (A) global differentiation is significant when compared to the (B) local differentiation. The figure was generated using the R package *adegenet;* Figure S7: Results of the spatial PCA analysis, lagged scores are plotted as a cline map, panel (A) representing first axis and (B) representing second axis. Black dots correspond sampling sites; Figure S8: Results of the (A) isolation by distance and (B) isolation by elevation plots illustrating the relationships between genetic differentiation among sites of wild *Agave angustifolia* var. *pacifica* from the state of Sonora, Mexico, and (A) geographic distance (r = 0.41, *p = 0.007*) and (B) isolation by elevation (r = 0.19, *p = 0.01*), using a two-dimensional kernel density estimation as implemented in *MASS* package in R; Figure S9: Boxplots of the individual based diversity estimates and inbreeding index for wild (green) and cultivated (blue) samples of *Agave angustifolia* var. *pacifica* from the state of Sonora, Mexico. No significant differences were found; Figure S10: Boxplots of the individual based diversity estimates and inbreeding index for wild *Agave angustifolia* var. *pacifica* from the state of Sonora, Mexico populations identified with spatial analysis. Significant differences between groups (after FDR correction) are represented by horizontal black line connecting groups that were significantly different, with asterisks corresponding to the significance level (*<0.05, **<0.001 and *** <0.0001); Figure S11: The overview diagram of relatedness coefficient as estimated with TrioML methodology among 95 agave individuals *Agave angustifolia* var. *pacifica* from the state of Sonora, Mexico. The degree of relatedness is represented by colors, from white that indicates no relationship, to red representing a clonal relationship. Samples are coded according to the Table S1.

## Author Contributions

Conceptualization, L.E.E. and A.K.; methodology, L.E.E., A.K., K.Y.R.M., F.M.F. and E.A.P.; software, bioinformatics analysis, and validation, A.K.; writing—original draft preparation, A.K.; writing—review and editing, L.E.E., A.K., K.Y.R.M., E.A.P. and F.M.F.; supervision, L.E.E.; project administration, L.E.E. E.A.P.; funding acquisition, L.E.E. All authors have read and agreed to the published version of the manuscript.

## Funding

This research was funded by PAPIIT project IG200122 provided to Dr. Luis E. Eguiarte and PAPIIT IN212321 provided to Dr. Daniel Piñero (Instituto de Ecología, UNAM), by National Autonomous University of Mexico, UNAM). Postdoctoral research fellowship for A.K. was provided by DGAPA/ UNAM IECO/S.ACAD/136/2021

## Institutional Review Board Statement

Not applicable.

## Informed Consent Statement

Not applicable.

## Data Availability Statement

The raw vcf file and additional stats file have been deposited in the ZENODO database with DOI: 10.5281/zenodo.6438381.

## Acknowledgments

We are thankful to Dr. Rafael Lira Saade (FES-Iztacala, UNAM) for his help in funding acquisition. We are also thankful for the Klimova A’s postdoctoral research fellowship provided by DGAPA/ UNAM IECO/S.ACAD/136/2021. We are thankful to Dr. Rosalinda Tapia and Dr. Marco Tulio Solano De la Cruz for specialized technical support in the laboratory and to the Q.B. José Fulgencio Martínez Rodríguez for crucial help in the field sampling, all of them from Instituto de Ecología, UNAM and Dr. Alejandra Moreno-Letelier (Instituto de Biología, UNAM) for the information on the standardization of GBS in other *Agave* species.

## Conflicts of Interest

The authors declare no conflict of interest.

## Notes

### Competing Interest Statement

The authors have declared no competing interest.

## References

1. Loveless, M.D.; Hamrick, J.L. Ecological determinants of genetic structure in plant populations. Annu. Rev. Ecol. S. 1984, 15(1), 65–95. https://doi.org/10.1146/annurev.es.15.110184.000433

2. Wright, S. The genetical structure of populations. Ann. Eugen. 1949, 15(1), 323–354. https://doi.org/10.1111/j.1469-1809.1949.tb02451.x

3. Pereira, H.M.; Navarro, L.M.; Martins, I.S. Global biodiversity change: the bad, the good, and the unknown. Annu. Rev. Environ. Resour. 2012, 37(1), 25–50. https://doi.org/10.1146/annurev-environ-042911-093511

4. Petit, R.J.; el Mousadik, A.; Pons, O. Identifying populations for conservation on the basis of genetic markers. Conserv. Biol. 1998, 12(4), 844–855. https://doi.org/10.1046/j.1523-1739.1998.96489.x

5. Hamrick, J.L.; Godt, M.J.W. Effects of life history traits on genetic diversity in plant species. Philos. T. R. Soc. Lond. B, Biol. Sci. 1996, 351(1345), 1291–1298. https://doi.org/10.1098/rstb.1996.0112

6. Nathan, R.; Muller-Landau, H.C. Spatial patterns of seed dispersal, their determinants and consequences for recruitment. Trends Ecol. Evol. 2000, 15(7), 278–285. https://doi.org/10.1016/s0169-5347(00)01874-7

7. Cruzan, M.B.; Hendrickson, E.C. Landscape genetics of plants: challenges and opportunities. Plant Commun. 2020, 1(6), 100100. https://doi.org/10.1016/j.xplc.2020.100100

8. Cain, M.L.; Milligan, B.G.; Strand, A.E. Long-distance seed dispersal in plant populations. Am. J. Bot. 2000, 87(9), 1217–1227. https://doi.org/10.2307/2656714

9. Nathan, R. Long-Distance dispersal of plants. Science 2006, 313(5788), 786–788. https://doi.org/10.1126/science.1124975

10. Bustamante, E.; Búrquez, A.; Scheinvar, E.; Eguiarte, L.E. Population genetic structure of a widespread bat-pollinated columnar cactus. PLOS ONE 2016, 11(3), e0152329. https://doi.org/10.1371/journal.pone.0152329

11. Cresswell, J.E.; Bassom, A.P.; Bell, S.A.; Collins, S.J.; Kelly, T.B. Predicted pollen dispersal by honey-bees and three species of bumble-bees foraging on oil-seed rape: a comparison of three models. Funct. Ecol. 1995, 9(6), 829. https://doi.org/10.2307/2389980

12. Hamrick, J.L.; Nason, J.D.; Fleming, T.H.; Nassar, J.M. Genetic diversity in columnar cacti. In Columnar cacti and their mutualists. Fleming, T.H., Valiente-Banuet, A., Eds.; The University of Arizona Press, Tucson, USA, 2002; pp. 122–133.

13. Robledo-Arnuncio, J.J.; Klein, E.K.; Muller-Landau, H.C.; Santamaría, L. Space, time and complexity in plant dispersal ecology. Mov. Ecol. 2014, 2(1). https://doi.org/10.1186/s40462-014-0016-3

14. Hamrick, J.L.; Murawski, D.A.; Nason, J.D. The influence of seed dispersal mechanisms on the genetic structure of tropical tree populations. Vegetatio 1993, 107–108(1), 281–297. https://doi.org/10.1007/bf00052230

15. Shay, J.E.; Pennington, L.K.; Montiel-Molina, J.A.; Toews, D.J.; Hendrickson, B.T.; Sexton, J.P. Rules of plant species ranges: applications for conservation strategies. Front. Ecol. Evol. 2021, 9. https://doi.org/10.3389/fevo.2021.700962

16. Arizaga, S.; Ezcurra, E.; Peters, E.; de Arellano, F.R.; Vega, E. Pollination ecology of Agave macroacantha (Agavaceae) in a Mexican tropical desert. II. The role of pollinators. Am. J. Bot. 2000, 87(7), 1011–1017. https://doi.org/10.2307/2657001

17. Duminil, J.; Fineschi, S.; Hampe, A.; Jordano, P.; Salvini, D.; Vendramin, G.; Petit, R. Can population genetic structure be predicted from life-history traits? Am. Nat. 2007, 169(5), 662–672. https://doi.org/10.1086/513490

18. Joshi, J.; Schmid, B.; Caldeira, M.C.; Dimitrakopoulos, P.G.; Good, J.; Harris, R.; Hector, A.; Huss-Danell, K.; Jumpponen, A.; Minns, A.; Mulder, C.P.H.; Pereira, J.S.; Prinz, A.; Scherer-Lorenzen, M.; Siamantziouras, A.S.D.; Terry, A.C.; Troumbis, A.Y.; Lawton, J.H. Local adaptation enhances performance of common plant species. Ecol. Lett. 2001, 4(6), 536–544. https://doi.org/10.1046/j.1461-0248.2001.00262.x

19. Nosil, P.; Funk, D.J.; Ortiz-Barrientos, D. Divergent selection and heterogeneous genomic divergence. Mol. Ecol. 2009, 18(3), 375–402. https://doi.org/10.1111/j.1365-294x.2008.03946.x

20. Orsini, L.; Vanoverbeke, J.; Swillen, I.; Mergeay, J.; de Meester, L. Drivers of population genetic differentiation in the wild: isolation by dispersal limitation, isolation by adaptation and isolation by colonization. Mol. Ecol. 2013, 22(24), 5983–5999. https://doi.org/10.1111/mec.12561

21. Brown, A.H.D. Human impact on plant gene pools and sampling for their conservation. Biol. Conserv. 1992, 62(2), 144. https://doi.org/10.1016/0006-3207(92)90950-r

22. Tilman, D.; Lehman, C. Human-caused environmental change: Impacts on plant diversity and evolution. Proc. Natl. Acad. Sci. 2001, 98(10), 5433–5440. https://doi.org/10.1073/pnas.091093198

23. Baucom, R.S.; Estill, J.C.; Cruzan, M.B. The effect of deforestation on the genetic diversity and structure in Acer saccharum (Marsh): Evidence for the loss and restructuring of genetic variation in a natural system. Conser. Genet. 2005, 6(1), 39–50. https://doi.org/10.1007/s10592-004-7718-9

24. Ahuja, M. R.; Jain, M.S. Genetic Diversity and Erosion in Plants: Case Histories, Softcover reprint of the original 1st 2016 ed.; Springer, 2019.

25. Khoury, C.K.; Brush, S.; Costich, D.E.; Curry, H.A.; Haan, S.; Engels, J.M.M.; Guarino, L.; Hoban, S.; Mercer, K.L.; Miller, A.J.; Nabhan, G.P.; Perales, H.R.; Richards, C.; Riggins, C.; Thormann, I. Crop genetic erosion: understanding and responding to loss of crop diversity. New Phytol. 2021, 233(1), 84–118. https://doi.org/10.1111/nph.17733

26. Reif, J.C.; Zhang, P.; Dreisigacker, S.; Warburton, M.L.; van Ginkel, M.; Hoisington, D.; Bohn, M.; Melchinger, A. E. Wheat genetic diversity trends during domestication and breeding. Theor. Appl. Genet. 2005, 110(5), 859–864. https://doi.org/10.1007/s00122-004-1881-8

27. van de Wouw, M.; van Hintum, T.; Kik, C.; van Treuren, R.; Visser, B. Genetic diversity trends in twentieth century crop cultivars: a meta analysis. Theor. Appl. Genet. 2010, 120(6), 1241–1252. https://doi.org/10.1007/s00122-009-1252-6

28. García-Mendoza, A. Los agaves de Mexico. Ciencias 2007, 087, 14-23. ISSN:0187-6376

29. Eguiarte, L.E.; Jiménez Barrón, O.A.; Aguirre-Planter, E.; Scheinvar, E.; Gámez, N.; Gasca-Pineda, J.; Castellanos-Morales, G.; Moreno-Letelier, A.; Souza, V. Evolutionary ecology of Agave: distribution patterns, phylogeny, and coevolution (an homage to Howard S. Gentry). Am. J. Bot. 2021, 108(2), 216–235. https://doi.org/10.1002/ajb2.1609

30. Slauson, L.A. Pollination biology of two chiropterophilous agaves in Arizona. Am. J. Bot. 2000, 87(6), 825– 836. https://doi.org/10.2307/2656890

31. Gentry, H.S. Agaves of Continental North America, 3rd ed.; University of Arizona Press, USA, 2004.

32. Good-Avila, S.V.; Souza, V.; Gaut, B.S.; Eguiarte, L.E. Timing and rate of speciation in Agave (Agavaceae). Proc. Natl. Acad. Sci. 2006, 103(24), 9124–9129. https://doi.org/10.1073/pnas.0603312103

33. Torres-García, I.; Rendón-Sandoval, F.J.; Blancas, J.; Casas, A.; Moreno-Calles, A.I. The genus Agave in agroforestry systems of Mexico. Bot. Sci. 2019, 97(3), 263. https://doi.org/10.17129/botsci.2202

34. Bowen, S.; Zapata, A.V. Geographical indications, terroir, and socioeconomic and ecological sustainability: The case of tequila. J. Rural Stud. 2009, 25(1), 108–119. https://doi.org/10.1016/j.jrurstud.2008.07.003

35. Bahre, C.J.; Bradbury, D.E. Manufacture of mescal in Sonora, Mexico. Econ. Bot. 1980, 34, 391–400. https://www.jstor.org/stable/4254220

36. Esqueda, M.; Coronado, M.L.; Gutiérrez, A.H.; Fragoso, T. Agave angustifolia Haw. Técnicas para el trasplante de vitroplantas a condiciones de agostadero. Secretaría de Agricultura Ganadería, Pesca y Alimentación (SAGARPA). México, 2013.

37. Council for Mescal Regulation. El Mezcal la cultura liquida de Mexico, 2020. https://comercam-dom.org.mx

38. Tetreault, D.; McCulligh, C.; Lucio, C. Distilling agro-extractivism: Agave and tequila production in Mexico. J. Agrar. Change. 2021, 21(2), 219–241. https://doi.org/10.1111/joac.12402

39. Luna Zamora, R. La construcción cultural y económico del tequila. Universidad de Guadalajara, Núcleo Universitario Los Belenes, 2015, ISBN: 978-607-8336-84-5.

40. Gil-Vega, K.; Díaz, C.; Nava-Cedillo, A.; Simpson, J. AFLP analysis of Agave tequilana varieties. Plant Sci. 2006, 170(4), 904–909. https://doi.org/10.1016/j.plantsci.2005.12.014

41. Vargas-Ponce, O.; Zizumbo-Villarreal, D.; Martínez-Castillo, J.; Coello-Coello, J.; Colunga-GarcíaMarín, P. Diversity and structure of landraces of Agave grown for spirits under traditional agriculture: A comparison with wild populations of A. angustifolia (Agavaceae) and commercial plantations of A. tequilana. Am. J. Bot. 2009, 96(2), 448–457. https://doi.org/10.3732/ajb.0800176

42. Dalton, R. Saving the agave. Nature 2005, 438(7071), 1070–1071. https://doi.org/10.1038/4381070a

43. Eguiarte, L.E.; Aguirre-Planter, E.; Aguirre, X.; Colín, R.; González, A.; Rocha, M.; Scheinvar, E.; Trejo, L.; Souza, V. From Isozymes to Genomics: population genetics and conservation of Agave in México. Bot. Rev. 2013, 79(4), 483–506. https://doi.org/10.1007/s12229-013-9123-x

44. Lindsay, D. L.; Swift, J.F.; Lance, R.F.; Edwards, C.E. A comparison of patterns of genetic structure in two co-occurring Agave species (Asparagaceae) that differ in the patchiness of their geographical distributions and cultivation histories. Bot. J. Linn. Soc. 2018, 186(3), 361–373. https://doi.org/10.1093/botlinnean/box099

45. Aguirre-Dugua, X.; Eguiarte, L.E. Genetic diversity, conservation and sustainable use of wild Agave cupreata and Agave potatorum extracted for mezcal production in Mexico. J. Arid Environ. 2013, 90, 36–44. https://doi.org/10.1016/j.jaridenv.2012.10.018

46. Figueredo, C.J.; Casas, A.; Colunga-GarcíaMarín, P.; Nassar, J. M.; González-Rodríguez, A. Morphological variation, management and domestication of ‘maguey alto’ (Agave inaequidens) and ‘maguey manso’ (A. hookeri) in Michoacán, México. J. Ethnobiol. Ethnomedicine. 2014, 10(1). https://doi.org/10.1186/1746-4269-10-66

47. Cabrera-Toledo, D.; Vargas-Ponce, O.; Ascencio-Ramírez, S.; Valadez-Sandoval, L.M.; Pérez-Alquicira, J.; Morales-Saavedra, J.; Huerta-Galván, O.F. Morphological and genetic variation in monocultures, forestry systems and wild populations of Agave maximiliana of Western Mexico: implications for its conservation. Front. Plant Sci. 2020, 11. https://doi.org/10.3389/fpls.2020.00817

48. Trejo, L.; Limones, V.; Peña, G.; Scheinvar, E.; Vargas-Ponce, O.; Zizumbo-Villarreal, D.; Colunga-GarcíaMarín, P. Genetic variation and relationships among agaves related to the production of Tequila and Mezcal in Jalisco. Ind. Crops Prod. 2018, 125, 140–149. https://doi.org/10.1016/j.indcrop.2018.08.072

49. Gentry, H.S. The Agave Family in Sonora. Agriculture Handbook no. 399. Agricultural Research Service, United States Department of Agriculture, Washington D.C., USA,1972.

50. Molina-Freaner, F.; Eguiarte, L.E. The pollination biology of two paniculate agaves (Agavaceae) from northwestern Mexico: contrasting roles of bats as pollinators. Am. J. Bot. 2003, 90(7), 1016–1024. https://doi.org/10.3732/ajb.90.7.1016

51. Menchaca, A.; Arteaga, M.C.; Medellin, R.A.; Jones, G. Conservation units and historical matrilineal structure in the tequila bat (Leptonycteris yerbabuenae). Glob. Ecol. Conserv. 2020, 23, e01164. https://doi.org/10.1016/j.gecco.2020.e01164

52. Wilkinson, G.S.; Fleming, T.H. Migration and evolution of lesser long-nosed bats Leptonycteris curasoae, inferred from mitochondrial DNA. Mol. Ecol. 1996, 5(3), 329–339. https://doi.org/10.1111/j.1365-294x.1996.tb00324.x

53. Morales-Garza, M.; Arizmendi, M.D.C.; Campos, J.; Martínez-Garcia, M.; Valiente-Banuet, A. Evidences on the migratory movements of the nectar-feeding bat Leptonycteris curasoae in Mexico using random amplified polymorphic DNA (RAPD). J. Arid Envir. 2007, 68(2), 248–259. https://doi.org/10.1016/j.jaridenv.2006.05.009

54. Westberg, E.; Kadereit, J.W. The influence of sea currents, past disruption of gene flow and species biology on the phylogeographical structure of coastal flowering plants. J. Biogeogr. 2009, 36(7), 1398–1410. https://doi.org/10.1111/j.1365-2699.2008.01973.x

55. de Witte, L.C.; Stöcklin, J. Longevity of clonal plants: why it matters and how to measure it. Ann. Bot. 2010, 106(6), 859–870. https://doi.org/10.1093/aob/mcq191

56. Arafeh, R.; Kadereit, J.W. Long-distance seed dispersal, clone longevity and lack of phylogeographical structure in the European distributional range of the coastal Calystegia soldanella (L.) R. Br. (Convolvulaceae). J. Biogeogr. 2006, 33(8), 1461–1469. https://doi.org/10.1111/j.1365-2699.2006.01512.x

57. Wieringa, J.G.; Boot, M.R.; Dantas-Queiroz, M.V.; Duckett, D.; Fonseca, E.M.; Glon, H.; Hamilton, N.; Kong, S.; Lanna, F.M.; Mattingly, K.Z.; Parsons, D.J.; Smith, M.L.; Stone, B.W.; Thompson, C., Zuo, L.; Carstens, B.C. Does habitat stability structure intraspecific genetic diversity? It’s complicated. Front. Biogeogr. 2020, 12(2). https://doi.org/10.21425/f5fbg45377

58. Silva-Arias, G.A.; Caballero-Villalobos, L.; Giudicelli, G.C.; Freitas, L.B. Landscape and climatic features drive genetic differentiation processes in a South American coastal plant. BMC Ecol. Evol. 2021, 21(1). https://doi.org/10.1186/s12862-021-01916-4

59. Savolainen, O.; Lascoux, M.; Merilä, J. Ecological genomics of local adaptation. Nat. Rev. Genet. 2013, 14(11), 807–820. https://doi.org/10.1038/nrg3522

60. Rodríguez-Garay, B.; Lomelí-Sención, J.; Tapia-Campos, E.; Gutiérrez-Mora, A.; García-Galindo, J.; Rodríguez-Domínguez, J.; Urbina-López, D.; Vicente-Ramírez, I. Morphological and molecular diversity of Agave tequilana Weber var. Azul and Agave angustifolia Haw. var. Lineño. Ind. Crops Prod. 2009, 29(1), 220– 228. https://doi.org/10.1016/j.indcrop.2008.05.007

61. Galinat, W.C. The domestication and genetic erosion of maize. Econ. Bot. 1974, 28, 31–37. https://doi.org/10.1007/BF02861376

62. Gaut, B.S.; Díez, C.M.; Morrell, P.L. Genomics and the contrasting dynamics of annual and perennial domestication. Trends Genet. 2015, 31(12), 709–719. doi:10.1016/j.tig.2015.10.002

63. Doebley, J.F.; Gaut, B.S.; Smith, B.D. The molecular genetics of crop domestication. Cell 2006, 127(7), 1309– 1321. https://doi.org/10.1016/j.cell.2006.12.006

64. Kantar, M.B.; Nashoba, A.R.; Anderson, J.E.; Blackman, B.K.; Rieseberg, L.H. The genetics and genomics of plant domestication. BioScience 2017, 67(11), 971–982. https://doi.org/10.1093/biosci/bix114

65. Smýkal, P.; Nelson, M.; Berger, J.; von Wettberg, E. The impact of genetic changes during crop domestication on healthy food development. Agronomy 2018, 8(3), 26. https://doi.org/10.3390/agronomy8030026

66. Liu, W.; Chen, L.; Zhang, S.; Hu, F.; Wang, Z.; Lyu, J.; Wang, B.; Xiang, H.; Zhao, R.; Tian, Z.; Ge, S.; Wang, W. Decrease of gene expression diversity during domestication of animals and plants. BMC Evol. Biol. 2019, 19(1). https://doi.org/10.1186/s12862-018-1340-9

67. Alfaro-Rojas, G.; Legaria, J.; Rodríguez-Pérez, J.; Genetic diversity in populations of pulquero agaves (agave spp.) in northeastern méxico state. Rev. Fitotec. Mex. 2007, 30(1), 1–12.

68. Figueredo-Urbina, C.J.; Álvarez-Ríos, G.D.; García-Montes, M.A.; Octavio-Aguilar, P. Morphological and genetic diversity of traditional varieties of agave in Hidalgo State, Mexico. PLOS ONE 2021, 16(7), e0254376. https://doi.org/10.1371/journal.pone.0254376

69. Rivera-Lugo, M.; García-Mendoza, A.; Simpson, J.; Solano, E.; Gil-Vega, K. Taxonomic implications of the morphological and genetic variation of cultivated and domesticated populations of the Agave angustifolia complex (Agavoideae, Asparagaceae) in Oaxaca, Mexico. Plant Syst. Evol. 2018, 304(8), 969–979. https://doi.org/10.1007/s00606-018-1525-0

70. Gross, B.L.; Olsen, K.M. Genetic perspectives on crop domestication. Trends Plant Sci. 2010, 15(9), 529–537. https://doi.org/10.1016/j.tplants.2010.05.008

71. Peláez, P.; Orona-Tamayo, D.; Montes-Hernández, S.; Valverde, M. E.; Paredes-López, O.; Cibrián-Jaramillo, A. Comparative transcriptome analysis of cultivated and wild seeds of Salvia hispanica (chia). Sci. Rep. 2019b, 9(1). https://doi.org/10.1038/s41598-019-45895-5

72. Yang, Z.; Li, G.; Tieman, D.; Zhu, G. Genomics Approaches to Domestication Studies of Horticultural Crops. Hortic. Plant J. 2019, 5(6), 240–246. https://doi.org/10.1016/j.hpj.2019.11.001

73. Liu, Y.; Du, H.; Li, P.; Shen, Y.; Peng, H.; Liu, S.; Zhou, G. A.; Zhang, H.; Liu, Z.; Shi, M.; Huang, X.; Li, Y.; Zhang, M.; Wang, Z.; Zhu, B.; Han, B.; Liang, C.; Tian, Z. Pan-Genome of Wild and Cultivated Soybeans. Cell 2020, 182(1), 162–176.e13. https://doi.org/10.1016/j.cell.2020.05.023

74. Bohra, A.; Kilian, B.; Sivasankar, S.; Caccamo, M.; Mba, C.; McCouch, S.R.; Varshney, R.K. Reap the crop wild relatives for breeding future crops. Trends Biotechnol. 2021, 40(4), 412–431. https://doi.org/10.1016/j.tibtech.2021.08.009

75. Félix-Valdez, L.I.; Vargas-Ponce, O.; Cabrera-Toledo, D.; Casas, A.; Cibrian-Jaramillo, A.; de la Cruz-Larios, L. Effects of traditional management for mescal production on the diversity and genetic structure of Agave potatorum (Asparagaceae) in central Mexico. Genet. Resour. Crop Evol. 2015, 63(7), 1255–1271. https://doi.org/10.1007/s10722-015-0315-6

76. Luna, R.; Epperson, B.K.; Oyama, K. High levels of genetic variability and inbreeding in two Neotropical dioecious palms with contrasting life histories. Heredity 2007, 99(4), 466–476. https://doi.org/10.1038/sj.hdy.6801027

77. Sonesson, A.K.; Meuwissen, T.H.E. Minimization of rate of inbreeding for small populations with overlapping generations. Genet. Res. 2001, 77(03). https://doi.org/10.1017/s0016672301005079

78. Trame, A.M.; Coddington, A.J.; Paige, K.N. Field and genetic studies testing optimal outcrossing in Agave schottii, a long-lived clonal plant. Oecologia 1995, 104(1), 93–100. https://doi.org/10.1007/bf00365567

79. Reed, D. H.; Frankham, R. Correlation between fitness and genetic diversity. Biol. Conserv. 2003, 17(1), 230– 237. https://doi.org/10.1046/j.1523-1739.2003.01236.x

80. Gómez-Ruiz, E.P.; Lacher Jr, T.E.; Moreno-Talamantes, A.; Flores Maldonado, J.J. Impacts of land cover change on the plant resources of an endangered pollinator. PeerJ 2021, 9, e11990. https://doi.org/10.7717/peerj.11990

81. McKey, D.; Elias, M.; Pujol, B.; Duputié, A. The evolutionary ecology of clonally propagated domesticated plants. New Phytol. 2010, 186(2), 318–332. https://doi.org/10.1111/j.1469-8137.2010.03210.x

82. Fisher, R.A. The genetical theory of natural selection, Clarendon Press, Oxford, UK, 1930. https://doi.org/10.5962/bhl.title.27468

83. Kondrashov, A.S. Deleterious mutations and the evolution of sexual reproduction. Nature 1988, 336(6198), 435–440. https://doi.org/10.1038/336435a0

84. National Research Council. Managing Global Genetic Resources: Agricultural Crop Issues and Policies. The National Academies Press, Washington, DC, USA, 1993. https://doi.org/10.17226/2116.

85. Jensen, E.L.; Díez-del-Molino, D.; Gilbert, M.T.P.; Bertola, L.D.; Borges, F.; Cubric-Curik, V.; de Navascués, M.; Frandsen, P.; Heuertz, M.; Hvilsom, C.; Jiménez-Mena, B.; Miettinen, A.; Moest, M.; Pečnerová, P.; Barnes, I.; Vernesi, C. Ancient and historical DNA in conservation policy. Trends Ecol. Evol. 2022. https://doi.org/10.1016/j.tree.2021.12.010

86. Moreno Salazar S.F.; Esqueda M.; Martínez J.; Palomino G. Tamaño del genoma y cariotipo en Agave angustifolia y A. rhodacantha de Sonora, México, Rev. Fitotec. Mex. 2007, 30(1).

87. García-Mendoza, A.; Chiang, F. The confusion of Agave vivipara L. and A. angustifolia Haw., two distinct taxa. Brittonia 2003, 55(1), 82–87. https://doi.org/10.1663/0007-196x

88. Yetman, D.; Van Devender, T.R. Mayo Ethnobotany: land, history, and traditional knowledge in northwestern Mexico. University of California Press, Berkeley, USA, 2002.

89. Bañuelos, N.; Salido, P. El mezcal en Sonora, México, más que una bebida espirituosa. Etnobotánica de Agave angustifolia Haw. Estudios Sociales 2012, 2, 173–197. https://www.redalyc.org/articulo.oa?id=41724972008

90. Salazar-Solano, V. La industria del bacanora: historia y tradición de resistencia en la sierra sonorense. Región y Sociedad 2007, 19(39), 105–133. https://doi.org/10.22198/rys.2007.39.a551

91. Doyle, J.J.; Doyle, J.L. A rapid DNA isolation procedure for small quantities of fresh leaf tissue. Phytochemistry 1987, 19, 11–15.

92. Vázquez-Lobo, A. Filogenia de hongos endófitos del género Pinus L.: Implementación de técnicas moleculares y resultados preliminares. Tesis de Licenciatura en Biología. Facultad de Ciencias, Universidad Nacional Autónoma de México, México, 1996.

93. Bolger, A.M.; Lohse, M.; Usadel, B. Trimmomatic: a flexible trimmer for Illumina sequence data. Bioinformatics 2014, 30(15), 2114–2120. https://doi.org/10.1093/bioinformatics/btu170

94. Eaton, D.A.R.; Overcast, I. ipyrad: Interactive assembly and analysis of RADseq datasets. Bioinformatics 2020, 36(8), 2592–2594. https://doi.org/10.1093/bioinformatics/btz966

95. Danecek, P.; Auton, A.; Abecasis, G.; Albers, C.A.; Banks, E.; DePristo, M.A.; Handsaker, R.E.; Lunter, G.; Marth, G.T.; Sherry, S.T.; McVean, G.; Durbin, R. The variant call format and VCFtools. Bioinformatics 2011, 27(15), 2156–2158. https://doi.org/10.1093/bioinformatics/btr330

96. O’Connor, T.D.; Fu, W.; Mychaleckyj, J.C.; Logsdon, B.; Auer, P.; Carlson, C.S.; Leal, S.M.; Smith, J.D.; Rieder, M.J.; Bamshad, M.J.; Nickerson, D.A.; Akey, J.M. Rare variation facilitates inferences of fine-scale population structure in humans. Mol. Biol. Evol. 2014, 32(3), 653–660. https://doi.org/10.1093/molbev/msu326

97. Pembleton, L.W.; Cogan, N.O.I.; Forster, J.W. StAMPP: an R package for calculation of genetic differentiation and structure of mixed-ploidy level populations. Mol. Ecol. Resour. 2013, 13(5), 946–952. https://doi.org/10.1111/1755-0998.12129

98. Alexander, D.H.; Novembre, J.; Lange, K. Fast model-based estimation of ancestry in unrelated individuals. Genome Res. 2009, 19(9), 1655–1664. https://doi.org/10.1101/gr.094052.109

99. Alexander, D.H.; Lange, K. Enhancements to the ADMIXTURE algorithm for individual ancestry estimation. BMC Bioinform. 2011, 12(1). https://doi.org/10.1186/1471-2105-12-246

100. Zheng, X.; Levine, D.; Shen, J.; Gogarten, S.M.; Laurie, C.; Weir, B.S. A high-performance computing toolset for relatedness and principal component analysis of SNP data. Bioinformatics 2012, 28(24), 3326–3328. https://doi.org/10.1093/bioinformatics/bts606

101. Caye, K.; Deist, T.M.; Martins, H.; Michel, O.; François, O. TESS3: fast inference of spatial population structure and genome scans for selection. Mol. Ecol. Resour. 2015, 16(2), 540–548. https://doi.org/10.1111/1755-0998.12471

102. Frichot, E.; Schoville, S.; Bouchard, G.; François, O. Correcting principal component maps for effects of spatial autocorrelation in population genetic data. Front. Genet. 2012, 3. https://doi.org/10.3389/fgene.2012.00254

103. Jombart, T. adegenet: a R package for the multivariate analysis of genetic markers. Bioinformatics 2008, 24(11), 1403–1405. https://doi.org/10.1093/bioinformatics/btn129

104. Thioulouse, J.; Chessel, D.; Champely, S. Multivariate analysis of spatial patterns: a unified approach to local and global structures. Environ. Ecol. Stat. 1995, 2(1), 1–14. https://doi.org/10.1007/bf00452928

105. Bougeard, S.; Dray, S. Supervised Multiblock Analysis in R with the ade4 Package. J. Stat. Softw. 2018, 86(1). https://doi.org/10.18637/jss.v086.i01

106. Ersts, P. Geographic distance matrix generatos (v. 1.2.3). American Museum of natural History, Center for Biodiversity and Conservation. Available online: https://biodiversityinformatics.amnh.org/open_source/gdmg/ (accessed on 1 March 2022)

107. Goudet, J. hierfstat, a package for r to compute and test hierarchical F-statistics. Mol. Ecol. Notes 2005, 5(1), 184–186. https://doi.org/10.1111/j.1471-8286.2004.00828.x

108. Alho, J.S.; Välimäki, K.; Merilä, J. Rhh: an R extension for estimating multilocus heterozygosity and heterozygosity–heterozygosity correlation. Mol. Ecol. Resour. 2010, 10(4), 720–722. https://doi.org/10.1111/j.1755-0998.2010.02830.x

109. Stoffel, M.A.; Esser, M.; Kardos, M.; Humble, E.; Nichols, H.; David, P.; Hoffman, J.I. inbreedR: an R package for the analysis of inbreeding. Methods Ecol. Evol. 2016, 7(11), 1331–1339. https://doi.org/10.1111/2041-210x.12588

110. Purcell, S.; Neale, B.; Todd-Brown, K.; Thomas, L.; Ferreira, M.A.; Bender, D.; Maller, J.; Sklar, P.; de Bakker, P.I.; Daly, M.J.; Sham, P.C. PLINK: A tool set for whole-genome association and population-based linkage analyses. Am. J. Hum. Genet. 2007, 81(3), 559–575. https://doi.org/10.1086/519795

111. Keller, M.C.; Visscher, P.M.; Goddard, M.E. Quantification of inbreeding due to distant ancestors and its detection using dense single nucleotide polymorphism data. Genetics 2011, 189(1), 237–249. https://doi.org/10.1534/genetics.111.130922

112. Yang, J.; Lee, S.H.; Goddard, M.E.; Visscher, P.M. GCTA: A tool for genome-wide complex trait analysis. Am. J. Hum. Genet. 2011, 88(1), 76–82. https://doi.org/10.1016/j.ajhg.2010.11.011

113. Marshall, T.C.; Coltman, D.; Pemberton, J.; Slate, J.; Spalton, J.A.; Guinness, F.E.; Smith, J.A.; Pilkington, J.G.; Clutton–Brock, T.H. Estimating the prevalence of inbreeding from incomplete pedigrees. Proc. Royal Soc. B. 2002, 269(1500), 1533–1539. https://doi.org/10.1098/rspb.2002.2035

114. R Core Team. R: A language and environment for statistical computing. R Foundation for Statistical Computing, Vienna, Austria. Available online https://www.R-project.org/ (accessed on 1 April 2022).

115. Wang, J. coancestry: a program for simulating, estimating and analysing relatedness and inbreeding coefficients. Mol. Ecol. Resour. 2010, 11(1), 141–145. https://doi.org/10.1111/j.1755-0998.2010.02885.x

116. Wang, J. Estimating pairwise relatedness in a small sample of individuals. Heredity 2017, 119(5), 302–313. https://doi.org/10.1038/hdy.2017.52

117. Kamvar, Z.N.; Tabima, J.F.; Grünwald, N.J. Poppr: an R package for genetic analysis of populations with clonal, partially clonal, and/or sexual reproduction. PeerJ 2014, 2, e281. https://doi.org/10.7717/peerj.281

