## Supplementary material for "Genomic analyses of wild and cultivated bacanora agave (*Agave angustifolia* var. *pacifica*) reveal inbreeding, few signs of cultivation history and shallow population structure"

This PDF file includes:

Tables S1 to S5

Figures S1 to S10

**Supplementary material tables.**

**Table S1.** Sites where individuals of *Agave angustifolia* var. *pacifica* from the state of Sonora were collected, including information on geographical region, site abbreviation, coordinates, number of individuals used for genomic analyses (*N*) and sample source (wild, cultivated, backyard plantation). Note: values of *N* do not include one individual removed during the genomic filtering step. All samples were collected in 2021.

| Id | Region | Site name | Latitude | Longitude | Source | N | Elevation, m |
|---|---|---|---|---|---|---|---|
| 1 | Bacanora | NovW1 | 28°57'37.1"N | 109°35'28.8"W | Wild | 3 | 576 |
| 2 | Bacanora | NovW2 | 28°56'51.5"N | 109°30'21.1"W | Wild | 3 | 899 |
| 3 | Bacanora | NovC1 | 28°59'43.2"N | 109°23'35.8"W | Cultivated | 3 | 503 |
| 4 | Bacanora | NovC2 | 28°58'42.5"N | 109°24'02.4"W | Backyard | 3 | 458 |
| 5 | Bacanora | NovC3 | 28°56'59.9"N | 109°30'53.9"W | Cultivated | 3 | 931 |
| 6 | Bacanora | NovW3 | 28°56'46.5"N | 109°40'10.7"W | Wild | 3 | 669 |
| 7 | Mazatan | MazC1 | 29°12'38.4"N | 110°07'20.2"W | Cultivated | 3 | 600 |
| 8 | Mazatan | MazW1 | 29°11'04.9"N | 110°09'00.3"W | Wild | 3 | 642 |
| 9 | Mazatan | MazW2 | 29°10'03.5"N | 110°09'17.0"W | Wild | 3 | 749 |
| 10 | Mazatan | MazC2 | 29°12'22.8"N | 110°08'34.9"W | Backyard | 3 | 591 |
| 11 | San Carlos | SCarW1 | 28°18'12.1"N | 111°24'02.4"W | Wild | 3 | 34 |
| 12 | San Carlos | SCarW2 | 27°58'37.4"N | 111°07'49.2"W | Wild | 3 | 7 |
| 13 | San Carlos | SCarW3 | 27°56'24.3"N | 111°05'51.4"W | Wild | 3 | 66 |
| 14 | Hermosillo | HermC1 | 29°00'49.7"N | 110°57'10.6"W | Cultivated | 3 | 276 |
| 15 | Hermosillo | HermC2 | 29°01'20.9"N | 110°57'00.2"W | Cultivated | 3 | 249 |
| 16 | Moctezuma | MocC1 | 29°27'34.5"N | 110°16'40.8"W | Cultivated | 3 | 422 |
| 17 | Moctezuma | MocW1 | 29°29'32.5"N | 110°11'55.8"W | Wild | 3 | 473 |
| 18 | Moctezuma | MocW2 | 29°46'15.4"N | 109°43'24.7"W | Wild | 3 | 807 |
| 19 | Moctezuma | MocW3 | 29°48'12.8"N | 109°38'20.7"W | Wild | 3 | 674 |
| 20 | Moctezuma | MocC2 | 29°41'08.3"N | 109°38'05.9"W | Cultivated | 3 | 615 |
| 21 | Moctezuma | MocC3 | 29°27'34.5"N | 110°16'40.8"W | Cultivated | 3 | 422 |
| 22 | Navojoa | NavW1 | 26°44'11.7"N | 109°34'09.5"W | Wild | 3 | 4 |
| 23 | Navojoa | NavW2 | 26°46'06.0"N | 109°28'47.2"W | Wild | 3 | 8 |
| 24 | Navojoa | NavW3 | 26°49'11.1"N | 109°17'18.5"W | Wild | 3 | 56 |
| 25 | Navojoa | NavC1 | 27°04'29.9"N | 109°20'41.4"W | Cultivated | 3 | 80 |
| 26 | Navojoa | NavC2 | 27°05'28.7"N | 109°06'18.9"W | Cultivated | 3 | 274 |
| 27 | Navojoa | NavC3 | 27°05'28.7"N | 109°06'18.9"W | Cultivated | 3 | 274 |
| 28 | Navojoa | NavC4 | 27°02'33.4"N | 108°58'50.8"W | Cultivated | 3 | 440 |

| | | | | | | | |
|---|---|---|---|---|---|---|---|
| 29 | Cam (Carretera Hermosillo-Guaymas) | CamW | 28°28'38.4"N | 111°02'43.7"W | Wild | 3 | 208 |
| 30 | Yecora | YecW1 | 28°49'14.2"N | 110°33'13.7"W | Wild | 2 | 447 |
| 31 | Yecora | YecW2 | 28°34'03.7"N | 109°41'37.6"W | Wild | 2 | 608 |
| 32 | Yecora | YecW3 | 28°34'21.3"N | 109°33'09.1"W | Wild | 1 | 188 |
| 33 | Yecora | YecW4 | 28°29'17.0"N | 109°22'09.7"W | Wild | 2 | 858 |
| 34 | Yecora | YecW5 | 28°25'57.6"N | 109°14'54.1"W | Wild | 2 | 645 |

**Table S2**. Quality information for the filtered SNPs used in the present study. The data shown is for 95 individuals of *Agave angustifolia* var. *pacifica* from the state of Sonora, Mexico, and 11, 619 SNPs.

| Quality statistic | Mean (SD) |
|---|---|
| Individual depth | 48.6 (6.5) |
| Individual missingness | 0.028 (0.01) |
| Site missingness | 0.028 (0.029) |

**Table S3.** Descriptive genomic diversity estimates and corresponding confidence intervals in parenthesis as determined for each management type. N - number of samples; MLH – multilocus heterozygosity; sMLH – standardized multilocus heterozygosity; $F_{IS}$ -Wright's inbreeding index; Fhat3 - inbreeding index; IR - internal relatedness.

| Management | N | MLH | sMLH | $F_{IS}$ | Fhat3 | IR |
|---|---|---|---|---|---|---|
| Wild | 53 | 0.22 (0.002) | 0.99 (0.013) | 0.13 (0.012) | 0.13 (0.012) | 0.05 (0.010) |
| Cultivated | 42 | 0.22 (0.003) | 1.00 (0.009) | 0.13 (0.007) | 0.13 (0.007) | 0.05 (0.006) |

**Table S4.** Descriptive genomic diversity statistics and corresponding confidence intervals in parenthesis for *Agave angustifolia* var. *pacifica* from the state of Sonora, Mexico based on population and management type. N - number of samples; MLH – multilocus heterozygosity; Fis -Wright's inbreeding index; Fhat3 - inbreeding index; IR - internal relatedness. Region/population column is coded accordingly to Table S1.

| Region/Population | N | Management | MLH | $F_{IS}$ | Fhat3 | IR |
|---|---|---|---|---|---|---|
| Novillo | 9 | Wild | 0.22 (0.015) | 0.15 (0.058) | 0.15 (0.066) | 0.07 (0.052) |
| Mazatan | 6 | Wild | 0.23 (0.011) | 0.09 (0.045) | 0.11 (0.041) | 0.04 (0.037) |
| San Carlos | 9 | Wild | 0.22 (0.002) | 0.14 (0.009) | 0.13 (0.010) | 0.06 (0.010) |
| Moctezuma | 8 | Wild | 0.22 (0.002) | 0.13 (0.009) | 0.13 (0.012) | 0.05 (0.009) |
| Navojoa | 9 | Wild | 0.22 (0.008) | 0.13 (0.319) | 0.11 (0.027) | 0.04 (0.029) |
| Camino (Cam), (Carretera Hermosillo-Guaymas) | 3 | Wild | 0.22 (0.015) | 0.15 (0.058) | 0.15 (0.033) | 0.08 (0.051) |
| Yecora | 9 | Wild | 0.22 (0.006) | 0.14 (0.025) | 0.13 (0.021) | 0.05 (0.023) |
| Hermosillo | 6 | Cultivated | 0.22 (0.010) | 0.15 (0.040) | 0.15 (0.046) | 0.06 (0.036) |
| Mazatan | 6 | Cultivated | 0.22 (0.008) | 0.12 (0.032) | 0.12 (0.016) | 0.04 (0.016) |
| Moctezuma | 9 | Cultivated | 0.22 (0.004) | 0.12 (0.016) | 0.13 (0.011) | 0.05 (0.010) |
| Navojoa | 12 | Cultivated | 0.22 (0.003) | 0.13 (0.012) | 0.13 (0.014) | 0.06 (0.008) |
| Novillo | 9 | Cultivated | 0.22 (0.003) | 0.12 (0.013) | 0.12 (0.014) | 0.04 (0.010) |

**Table S5.** Number of loci and raw SNPs recovered for the datasets under different ipyrad clustering parameters

| Samples | Clustering parameter | Loci | SNPs |
|---|---|---|---|
| All sites | 0.85 | 86,014 | 610,119 |
| | 0.9 | 94, 174 | 638,704 |
| | 0.95 | 111,390 | 598,158 |

**Supplementary material figures.**

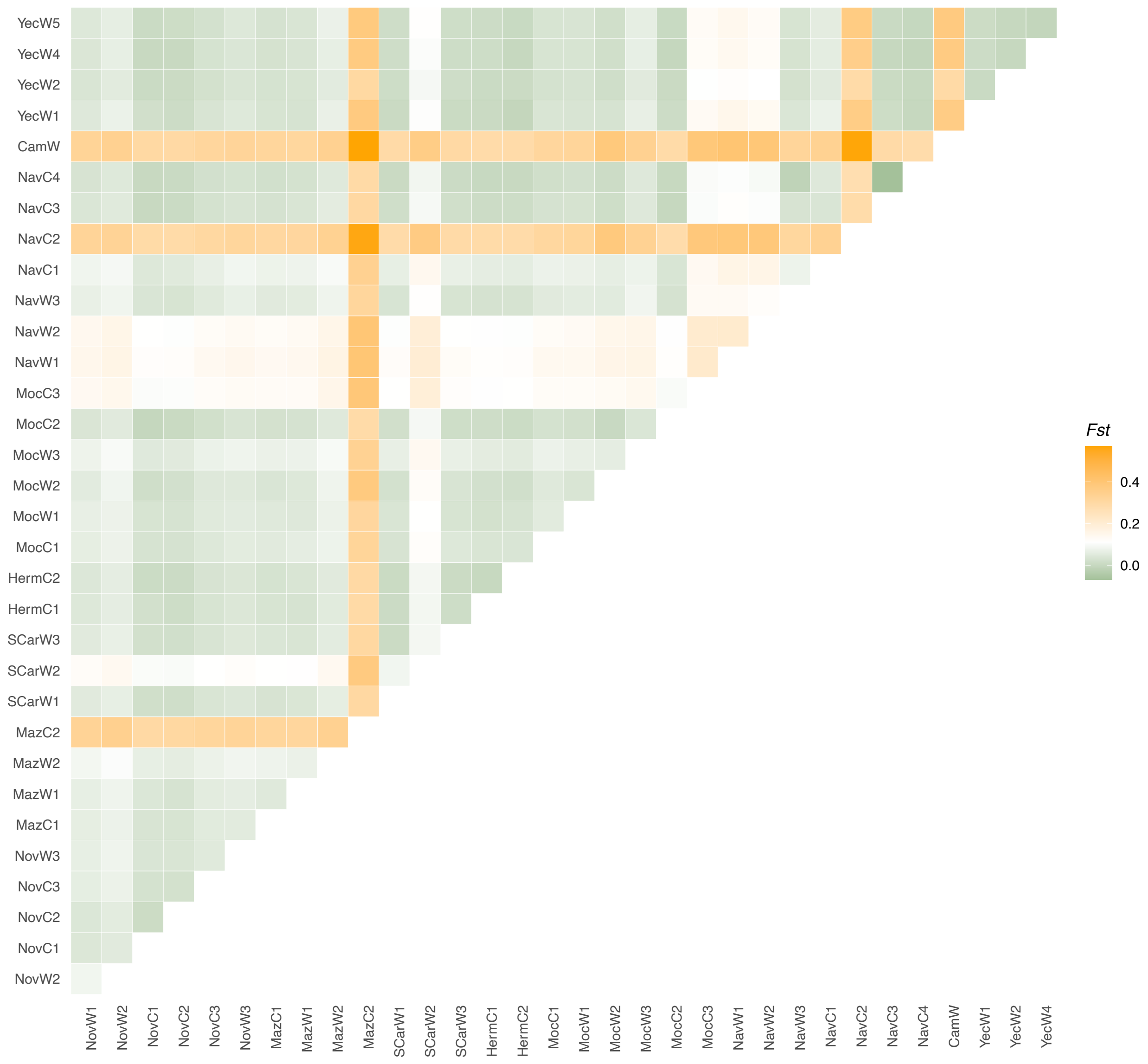


**Figure S1**. Pairwise $F_{ST}$ differences among all the sampling sites of wild and cultivated *Agave angustifolia* var. *pacifica* from the state of Sonora, Mexico. Colors represent $F_{ST}$ values from the lowest of -0.07 in green to the highest of 0.57 in orange. Sampling sites are coded accordingly to Table S1.

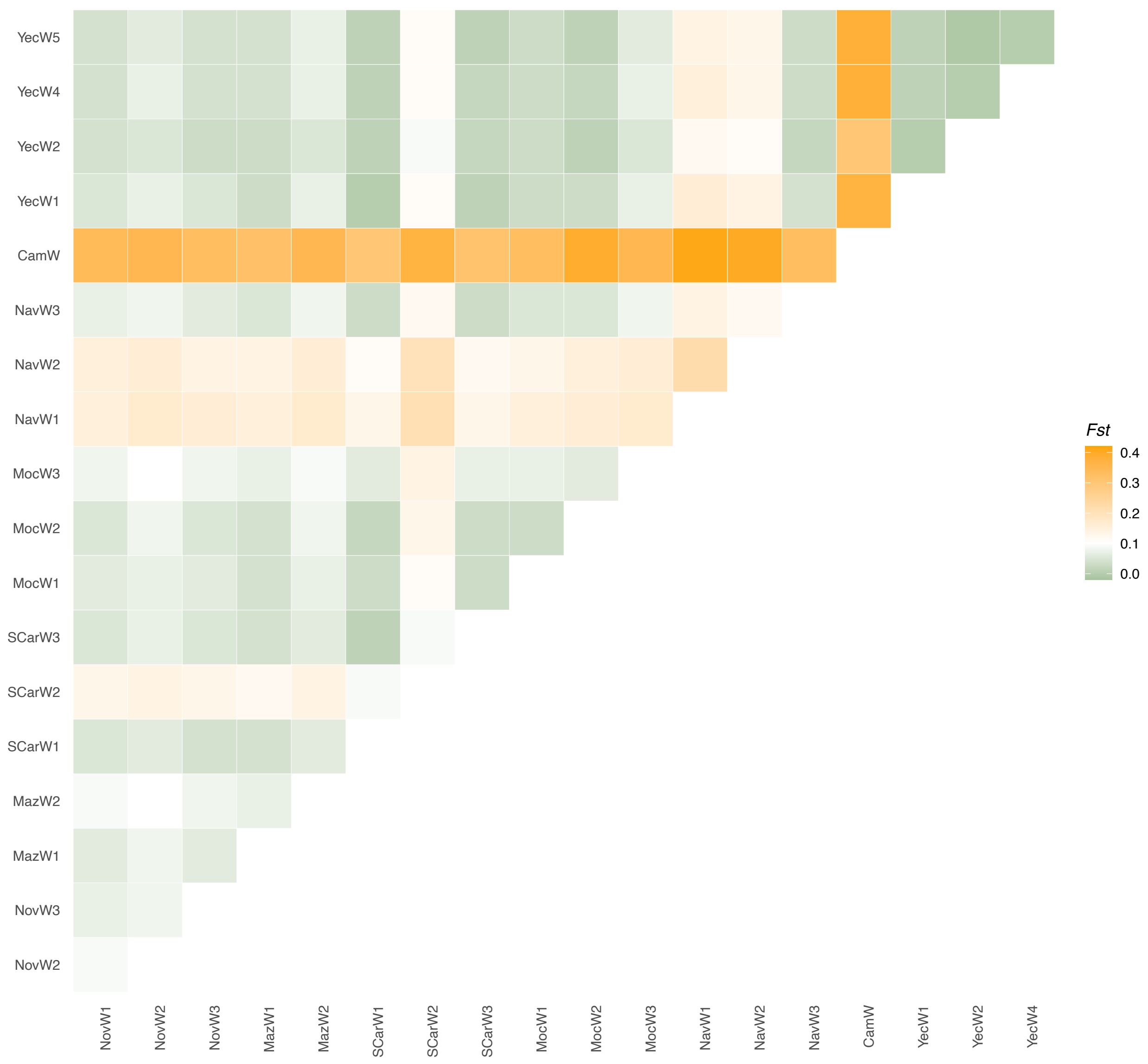


**Figure S2**. Pairwise $F_{ST}$ differences among all the sampling sites of wild *Agave angustifolia* var. *pacifica* from the state of Sonora, Mexico. Colors represent $F_{ST}$ values from the lowest of -0.01 in green to the highest of 0.41 in orange. Sampling sites are coded accordingly to Table S1.

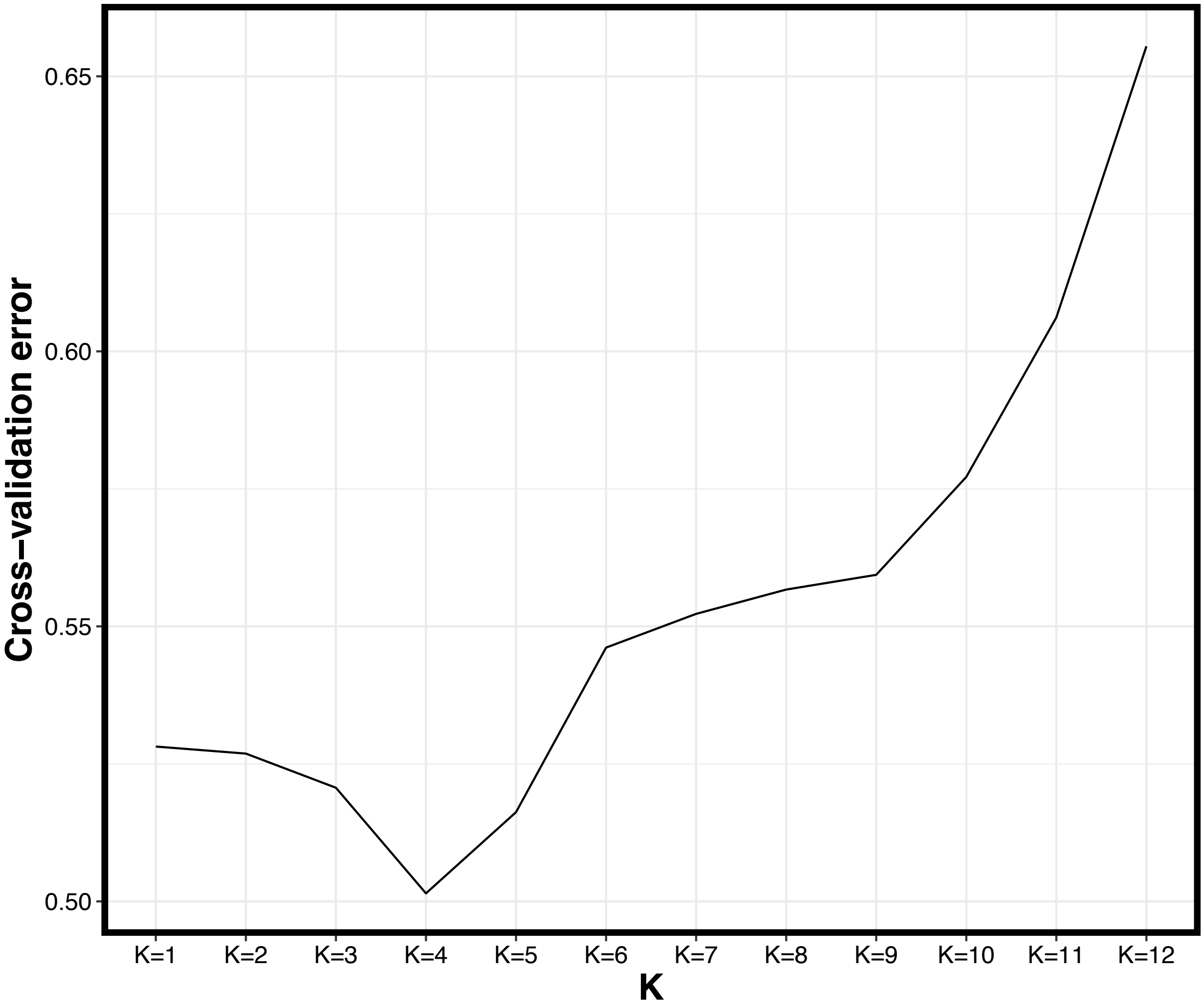


**Figure S3.** Plot of ADMIXTURE cross validation error from *K*=1 through *K*=12, as based on all samples (cultivated and wild).

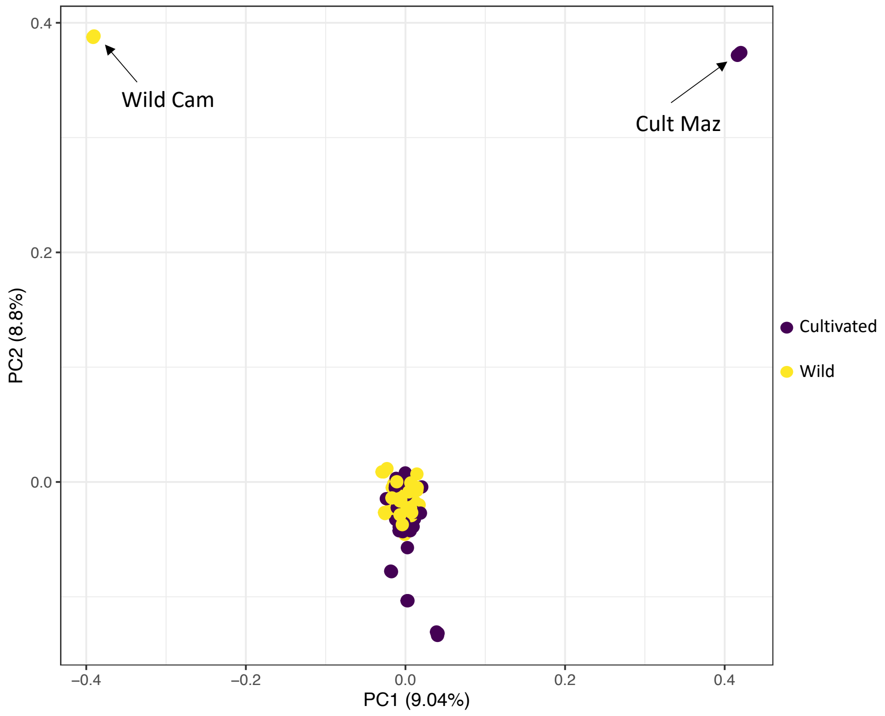


**Figure S4.** Relationships among *Agave angustifolia* var. *pacifica* individuals from the state of Sonora, Mexico, wild and cultivated samples as represented by principal component analysis (PCA) using 11,619 genome-wide SNPs. All 95 samples were included.

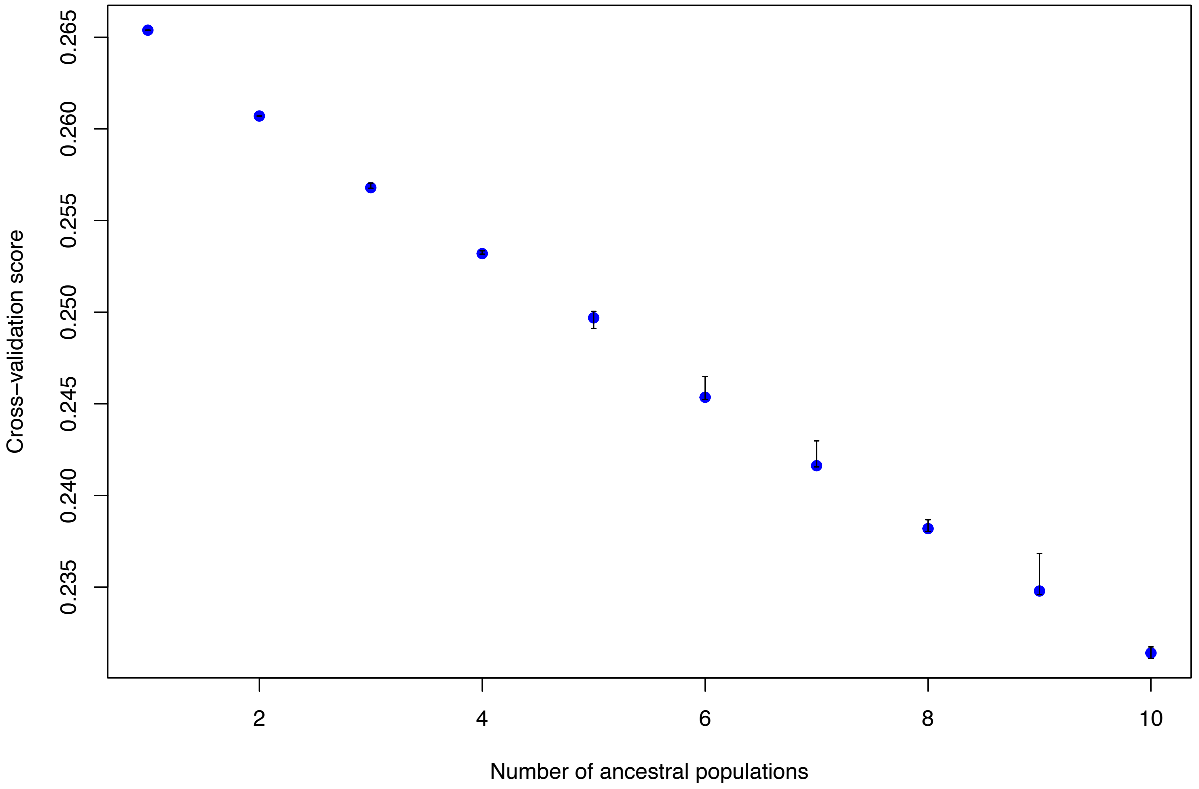


**Figure S5.** Plot of TESS3 cross validation error from *K*=1 through *K*=10, based only on wild samples. Twenty runs were performed for each value of *K*.

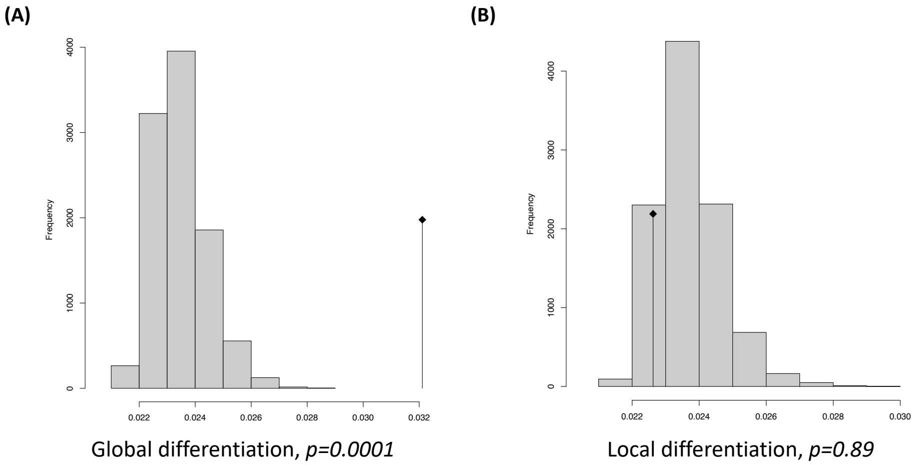


**Figure S6.** Results of Eigen value test for sPCA analysis. The (A) global differentiation is signif icant when compared to the (B) local differentiation. The figure was generated using the R packa ge *adegenet*.

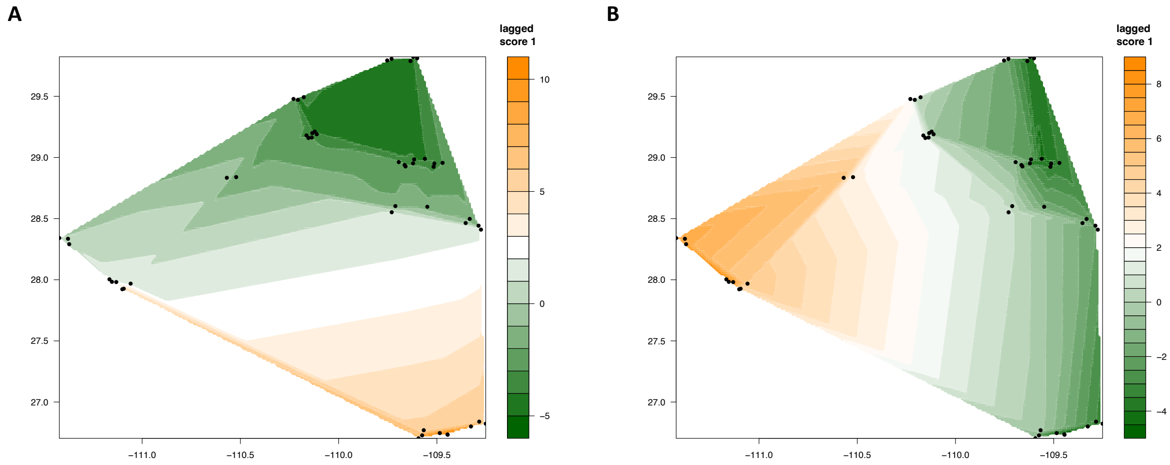


**Figure S7.** Results of the spatial PCA analysis, lagged scores are plotted as a cline map, panel (A ) representing first axis and (B) representing second axis. Black dots correspond sampling sites.

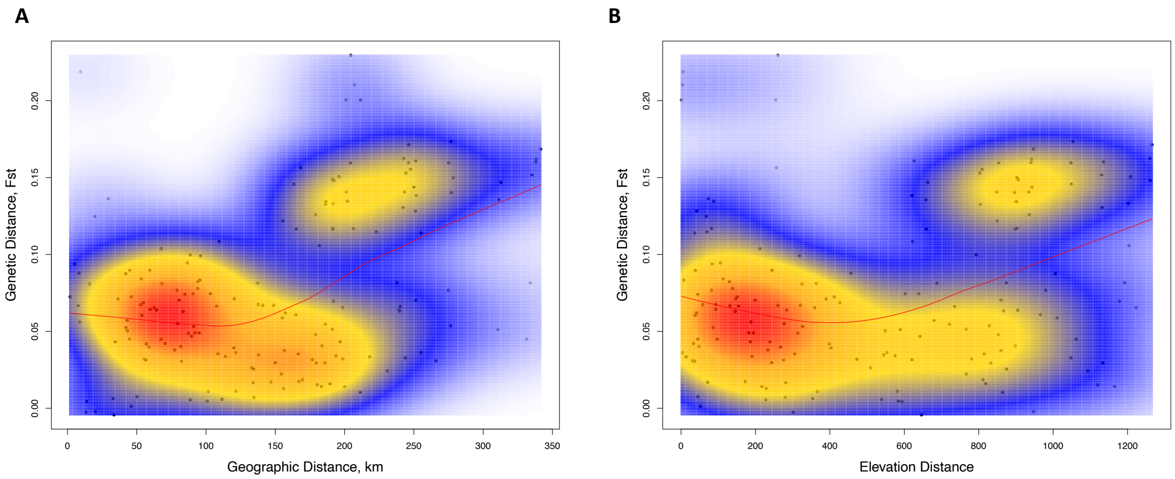


**Figure S8.** Results of the (A) isolation by distance and (B) isolation by elevation plots illustrating the relationships between genetic differentiation among sites of wild *Agave angustifolia* var. *pacifica* from the state of Sonora, Mexico, and (A) geographic distance (r = 0.41, $p = 0.007$) and (B) isolation by elevation (r = 0.19, $p = 0.01$), using a two-dimensional kernel density estimation as implemented in *MASS* package in R.

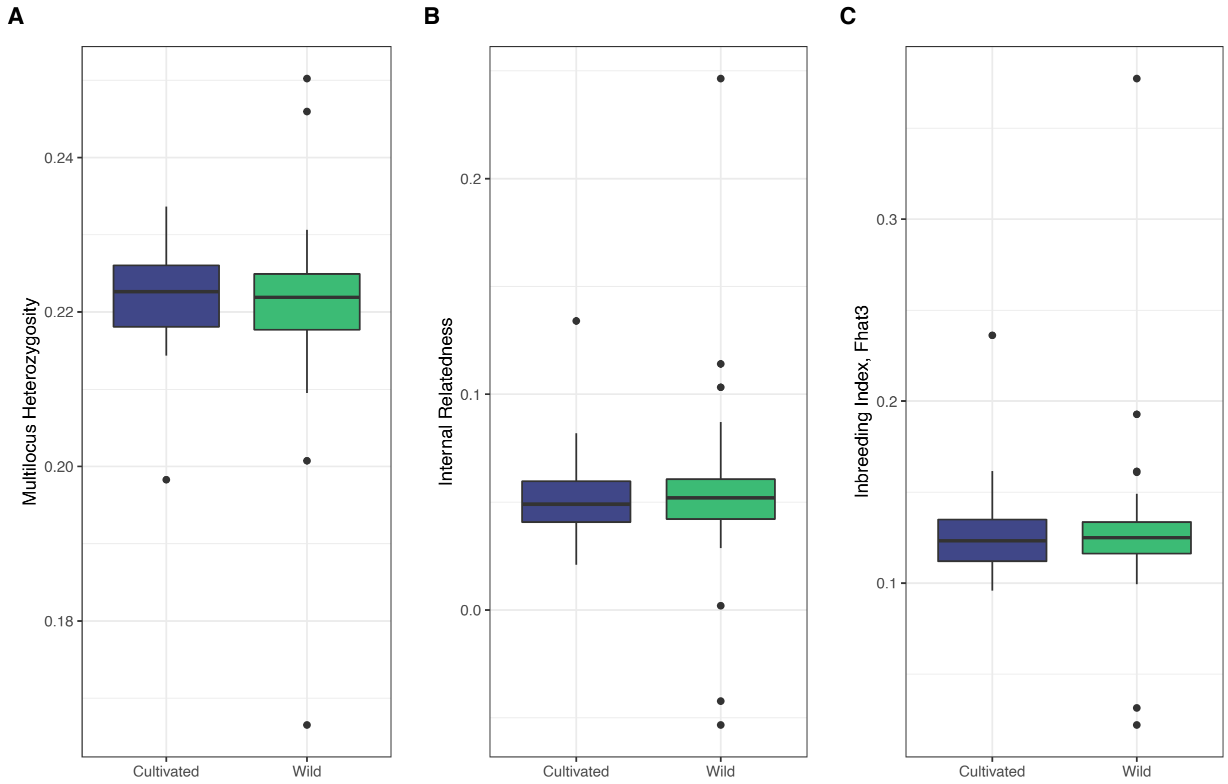


**Figure S9.** Boxplots of the individual based diversity estimates and inbreeding index for wild (green) and cultivated (blue) samples of *Agave angustifolia* var. *pacifica* from the state of Sonora, Mexico. No significant differences were found.

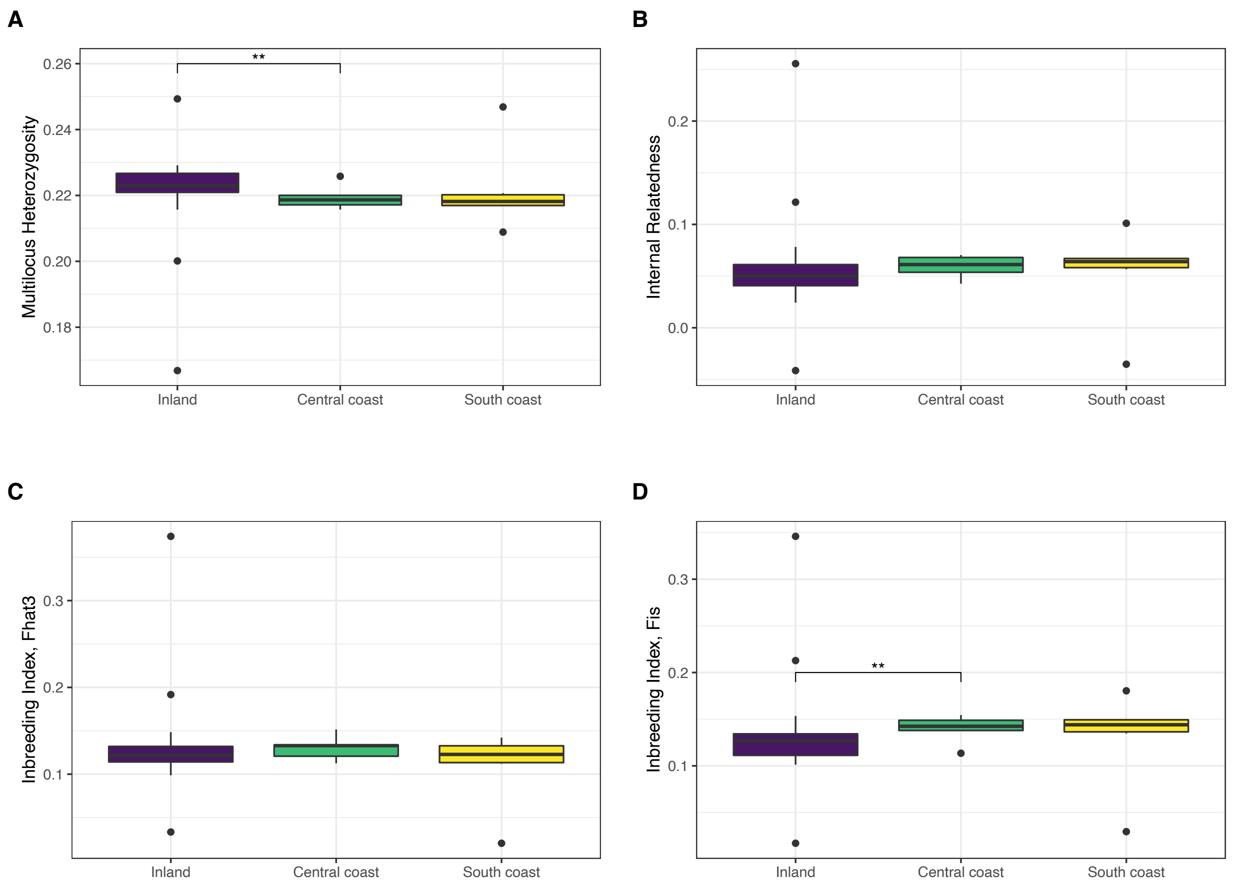


**Figure S10.** Boxplots of the individual based diversity estimates and inbreeding index for wild *Agave angustifolia* var. *pacifica* from the state of Sonora, Mexico populations identified with spatial analysis. Significant differences between groups (after FDR correction) are represented by horizontal black line connecting groups that were significantly different, with asterisks corresponding to the significance level (*<0.05, **<0.001 and *** <0.0001).
